# Environmentally robust *cis*-regulatory changes underlie rapid climatic adaptation

**DOI:** 10.1101/2022.08.29.505745

**Authors:** Mallory A. Ballinger, Katya L. Mack, Sylvia M. Durkin, Eric A. Riddell, Michael W. Nachman

## Abstract

Changes in gene expression are proposed to play a major role in adaptive evolution. While it is known that gene expression is highly sensitive to the environment, very few studies have determined the influence of genetic and environmental effects on adaptive gene regulation in natural populations. Here, we utilize allele-specific expression to characterize *cis* and *trans* gene regulatory divergence in temperate and tropical house mice in two metabolic tissues under two thermal conditions. First, we show that gene expression divergence is pervasive between populations and across thermal conditions, with roughly 5-10% of genes exhibiting genotype-by-environment interactions. Second, we found that most expression divergence was due to *cis*-regulatory changes that were stable across temperatures. In contrast, patterns of expression plasticity were largely attributable to *trans*-effects, which showed greater sensitivity to temperature. Nonetheless, we discovered a small subset of temperature-dependent *cis*-regulatory changes, thereby identifying loci underlying expression plasticity. Finally, we performed scans for selection in wild house mice to identify genomic signatures of rapid adaptation. Genomic outliers were enriched in genes with evidence for *cis*-regulatory divergence. Notably, these genes were associated with phenotypes that affected body weight and metabolism, suggesting that *cis*-regulatory changes are a possible mechanism for adaptive body size evolution between populations. Our results show that gene expression plasticity, largely controlled in *trans*, may facilitate the colonization of new environments, but that evolved changes in gene expression are largely controlled in *cis*, illustrating the genetic and non-genetic mechanisms underlying the establishment of populations in new environments.

**Significance Statement:** Gene expression variation is shaped by both genetic and environmental effects, yet these two factors are rarely considered together in the context of adaptive evolution. We studied environmental influences on gene regulatory evolution in temperate and tropical house mice in cold and warm laboratory environments. We discovered that genetic effects in the form of *cis*-regulatory divergence were pervasive and largely insensitive to the environment. Many of these genetic effects are under selection and are associated with genes that affect body size, suggesting *cis*-regulatory changes as a possible mechanism for adaptive body size evolution. We also discovered many *trans*-effects controlling expression plasticity, demonstrating the importance of both genetic and non-genetic changes associated with adaptation over short timescales (a few hundred generations).

---

A major goal in evolutionary biology is to understand how organisms adapt to novel environments. Changes in gene expression have long been recognized to play a major role in adaptive evolution (1, 2), especially across short evolutionary timescales (3, 4). Gene expression is highly sensitive to the environment and genotype-by-environment interactions (GxE) constitute a large proportion of gene expression variation (5–8). Moreover, selection on genetic variation underlying plasticity may facilitate the colonization of new environments, especially during the initial stages of adaptation (9–11). Yet, we have a relatively poor understanding of how expression plasticity is controlled and how regulatory architecture evolves in populations adapting to different environments. For instance, while numerous studies have supported the evolution of gene expression through *cis*-regulatory changes (e.g., mutations in promoters, enhancers) (12–16), the extent to which these changes are environmentally sensitive and modulate expression plasticity is not well understood. *Trans*-effects (e.g., transcription factors) may also play a significant role in plastic changes in gene expression by modifying gene regulatory networks. Selection may then favor divergence through *trans*acting mechanisms when such changes are beneficial in new environments (17, 18). Determining how both genetic and environmental effects influence the evolution of gene expression differences in natural populations is key to understanding the molecular mechanisms of adaptation.

The recent expansion of house mice into the Americas provides an opportunity to address the environmental sensitivity of gene regulatory changes involved in adaptive evolution. Since their arrival from Western Europe ∼500 years ago, house mice (*Mus musculus domesticus*) have rapidly adapted to various climatic extremes through changes in morphology, physiology, and behavior (19–23). One striking example of this is changes in body size, as mice from more northern populations are significantly larger than mice closer to the equator, likely reflecting adaptation to differing thermal environments (23). Previous studies point to an important role for gene regulation in driving this local adaptation. First, genomic scans have primarily identified positive selection on noncoding regions (20, 21), which have been linked to differences in gene expression (21, 24). Second, changes in *cis*-regulation at specific loci have been associated with variation in body weight in North American mice (24). Finally, gene expression plasticity has been shown to differ between populations in response to environmental stressors (22), suggesting a role for environment-specific regulatory divergence in local adaptation.

Here, we investigate the role of gene regulation in the rapid adaptation of house mice to contrasting thermal environments. Specifically, using RNA-seq data collected from liver and brown adipose tissue in males and females, we measured gene expression divergence in inbred lines of temperate and tropical mice and in their F1 hybrids when reared under warm and cold temperatures. This allowed us to describe the proportion of divergently expressed genes that are due to changes in *cis, trans*, or both, and to determine the degree to which *cis*- and *trans*-regulation is temperature-dependent. Finally, we performed scans for selection in wild populations of house mice to identify genomic signatures of adaptation. We then intersected these genomic outliers with genes exhibiting *cis*-regulatory divergence to identify *cis*-regulatory mutations associated with local adaptation. Our results provide insight into how gene regulation changes in response to the environment and how complex regulatory divergence within a species may contribute to adaptive evolution.

## Results

### Extensive gene expression divergence between temperate and tropical house mice

To characterize the regulatory architecture of adaptation, we first examined gene expression differences in two wild-derived inbred lines of house mice from different environments in the Americas: Saratoga Springs, New York, USA (SARA), located at 43^*°*^N, and Manaus, Amazonas, Brazil (MANA), located near the equator at 3^*°*^S. Saratoga Springs and Manaus differ considerably in climate, such as mean annual temperature (Fig. 1A), and mice from these environments show population-level differences in various traits, including morphology and gene expression (21, 23) (*SI Appendix, Supplementary Text*). Specifically, mice from New York are larger, retain more heat through their fur, and have shortened extremities compared to mice from Brazil (ANOVA tests, *P* < 0.05) (Fig. 1B and *SI Appendix*, Table S1), suggesting adaptation to different climates (23).

**Fig. 1.**
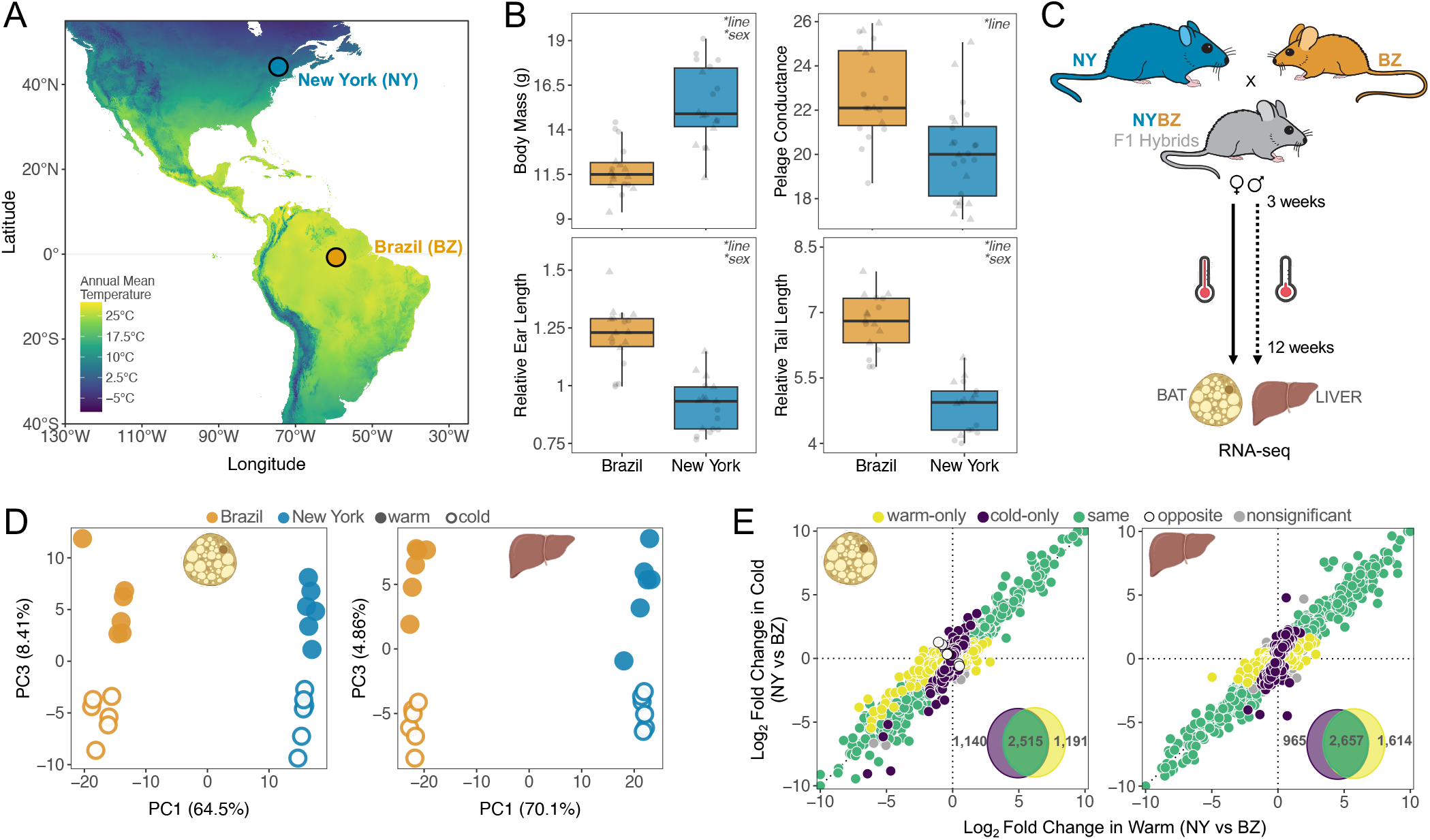
Evolved differences in phenotypes and gene expression. **(A)** Variation in mean annual temperature across North and South America. Wild-derived inbred lines originate from upstate New York (43^*°*^N) and equatorial Brazil (3^*°*^S). **(B)** Differences in body mass (g), pelage conductance (W m^-2 ∼^C^-1^), tail length (mm), and ear length (mm) between wild-derived inbred lines of New York (SARA) and Brazil (MANA). Tail length and ear length are plotted relative to body mass for each individual. Individuals are represented as individual points, and boxplots indicate the 25th, median, and 75th quartiles. Results from linear mixed models are presented in upper right corners (*\*P* < 0.05; *SI Appendix*, Table S1). Males (circles) and females (triangles) show similar patterns and are combined for plotting simplicity. **(C)** Common garden experimental design. Individuals were reared under two temperatures from weaning until adults. **(D)** Principal component plots for PC1 vs PC3 based on male gene expression in BAT and liver. PC1 separates individuals based on genotype while PC3 reflects environmental differences. Principle component plots for PC1 vs PC2 are provided in *SI Appendix*, Figs. S1-S2. **(E)** Expression divergence between New York and Brazil males in warm and cold for both BAT and liver. Log2 fold changes between parents were calculated for all genes independently. In each panel, points (representing individual genes) are colored depending on their direction and significance of the log2 fold change. Insets depict the total number of differentially expressed genes for each comparison (FDR < 0.05). Females show similar patterns and are depicted in *SI Appendix*, Figs. S2-S3.

We explored patterns of gene expression evolution by rearing inbred lines from New York and Brazil under two temperatures (5^*°*^C and 21^*°*^C) and sequenced brown adipose tissue (BAT) and liver transcriptomes of 48 individuals (6 / line / sex / environment) (Fig. 1C). We chose these two tissues as they play important roles in both metabolism and adaptive thermogenesis (25–27). Principal component analysis (PCA) of all gene expression data revealed tissue type as the largest source of variance (PC1 ∼97% of variance explained), followed by sex (PC2 ∼1.5%) (*SI Appendix*, Fig. S1). Within each tissue and sex, New York and Brazil mice cleanly separated along PC1 (>60% of variance explained), while PC3 largely separated warm- and cold-reared mice (>4% of variance explained) (Fig. 1D and *SI Appendix*, Fig. S2). We also identified more than a third of genes to be differentially expressed between New York and Brazil mice (false discovery rate (FDR) < 0.05) (Fig. 1E and *SI Appendix*, Figs. S3-S4), with most expression differences concordant across environments and sexes.

This strong pattern of divergence between lines was also apparent when we categorized differentially expressed genes as those showing genetic variation (G), environmental variation [i.e., plasticity (E)], or genetic variation for plasticity (i.e., GxE) (Fig. 2A and *SI Appendix*, Fig. S3). Genotype had >1.5x larger effect size (calculated as the mean absolute value of the log2 fold change) on differential gene expression than environment across both tissues (Fig. 2A and *SI Appendix*, Fig. S3). Similar effects were identified when we attributed expression differences to genotype and sex, though these patterns were largely tissue-dependent (*SI Appendix*, Fig. S4). Overall, these results demonstrate that within sexes and tissues, genotype plays a larger role than either environment or GxE interactions in shaping expression differences between temperate and tropical house mice.

**Fig. 2.**
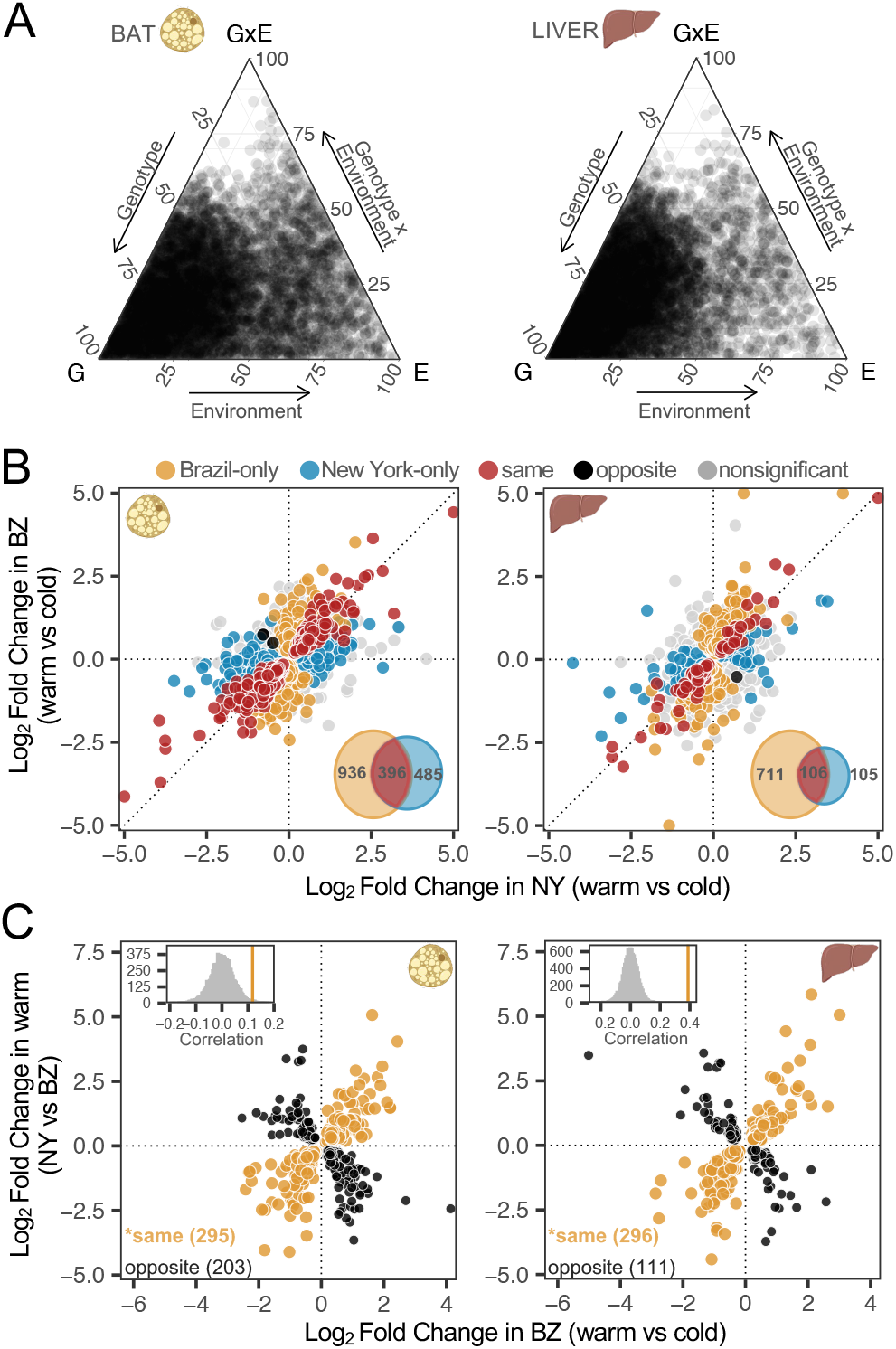
Patterns of genotype-by-environment interactions (GxE) **(A)** Ternary plots depicting the proportion of each gene’s expression variance explained by genotype (G), environment (E), and GxE. The relative proportion of each factor is shown for all differentially expressed male genes in BAT and liver. Total variance is the sum of all three components. **(B)** Comparison of gene expression differences between temperature regimes in NY and BZ males in both tissues. Log2 fold changes between temperatures were calculated for all genes independently. In each panel, points (representing individual genes) are colored depending on their direction and significance of the log2 fold change. GxE categories include line-specific responses or opposite responses between lines (see Methods). Insets depict the total number of differentially expressed genes for each comparison (FDR < 0.05). **(C)** The relationship between gene expression plasticity and evolved divergence in both tissues. Points represent expression differences with statistically significant plasticity in BZ (cold vs warm; FDR < 0.05) as well as significant expression divergence between NY and BZ at warm temperature (FDR < 0.05). Points colored in orange represent genes with a positive correlation between plasticity and evolved divergence, while points in black represent genes with a negative correlation. Insets depict the observed correlation coefficient (orange solid lines) is more positive than a randomized distribution of correlation coefficients for each tissue (see Methods). Asterisks denote significance of plasticity for each tissue (binomial exact tests, *P* < 0.05). Females show similar patterns and are depicted in *SI Appendix*, Figs. S2-S3.

### Reduced gene expression plasticity in cold-adapted mice

Given that New York and Brazil mice have evolved under different thermal environments, we reasoned that gene expression responses to temperature would differ between these lines. Roughly ∼5% and ∼10% of all expressed genes showed significant GxE in liver and BAT, respectively (FDR < 0.05) (Fig. 2B and *SI Appendix*, Fig. S3 and Table S2). Although 3 genes showed opposite responses between lines across BAT (*cyfip2, wnt11*) and liver (*cmpk2*), most GxE patterns were categorized as line-specific in both tissues. Notably, we found fewer differentially expressed genes between environmental conditions in New York mice (∼5% BAT; ∼1% liver) than Brazil mice (∼10% BAT; ∼5% liver) (Chi-square tests, liver and BAT: *P* < 0.05), suggesting that New York mice may be less sensitive to cold stress.

Next, we explored the relationship between gene expression plasticity and evolved gene expression differences. Plasticity may facilitate the colonization of new environments by moving a population closer to the phenotypic optimum, or alternatively, reduce fitness under new environmental stressors (10, 28). To determine if the pronounced transcriptional response to temperature of Brazil mice aligns with expression divergence between lines (*sensu* refs. 29, 30), we asked whether the direction of expression plasticity of Brazil mice correlates with total expression divergence between New York and Brazil mice (see Methods). We found that expression plasticity generally goes in the same direction as evolved divergence (positive Spearman’s correlations in both tissues, *P* < 0.05) (Fig. 2C and *SI Appendix*, Fig. S3), consistent with the idea that this plasticity is adaptive (22, 30–32). However, this pattern was less prominent in BAT, with only slightly more genes exhibiting concordant than discordant expression patterns (Fig. 2C).

### Expression divergence is predominantly due to *cis*-regulatory changes, which are enriched for body size and metabolism

To investigate the gene regulatory mechanisms underlying expression differences between New York and Brazil mice, we generated BAT and liver RNA-seq from NY x BZ F1 hybrids reared in both warm and cold environments (Fig. 1C and *SI Appendix*, Fig. S5). Measuring gene expression in F1 hybrids allowed us to discern if parental gene expression differences are due to *cis*- and/or *trans*-acting changes by assessing patterns of allele-specific expression (ASE) (Fig. 3A). Specifically, as F1 hybrids inherit both a Brazil allele and New York allele within the same *trans*-acting environment, differences in expression between alleles are indicative of one or more *cis*-acting elements (33–36). In contrast, if no ASE is detected in hybrids but differences are observed between parental lines, we can infer divergence is likely due to *trans*-acting factors (33–36).

**Fig. 3.**
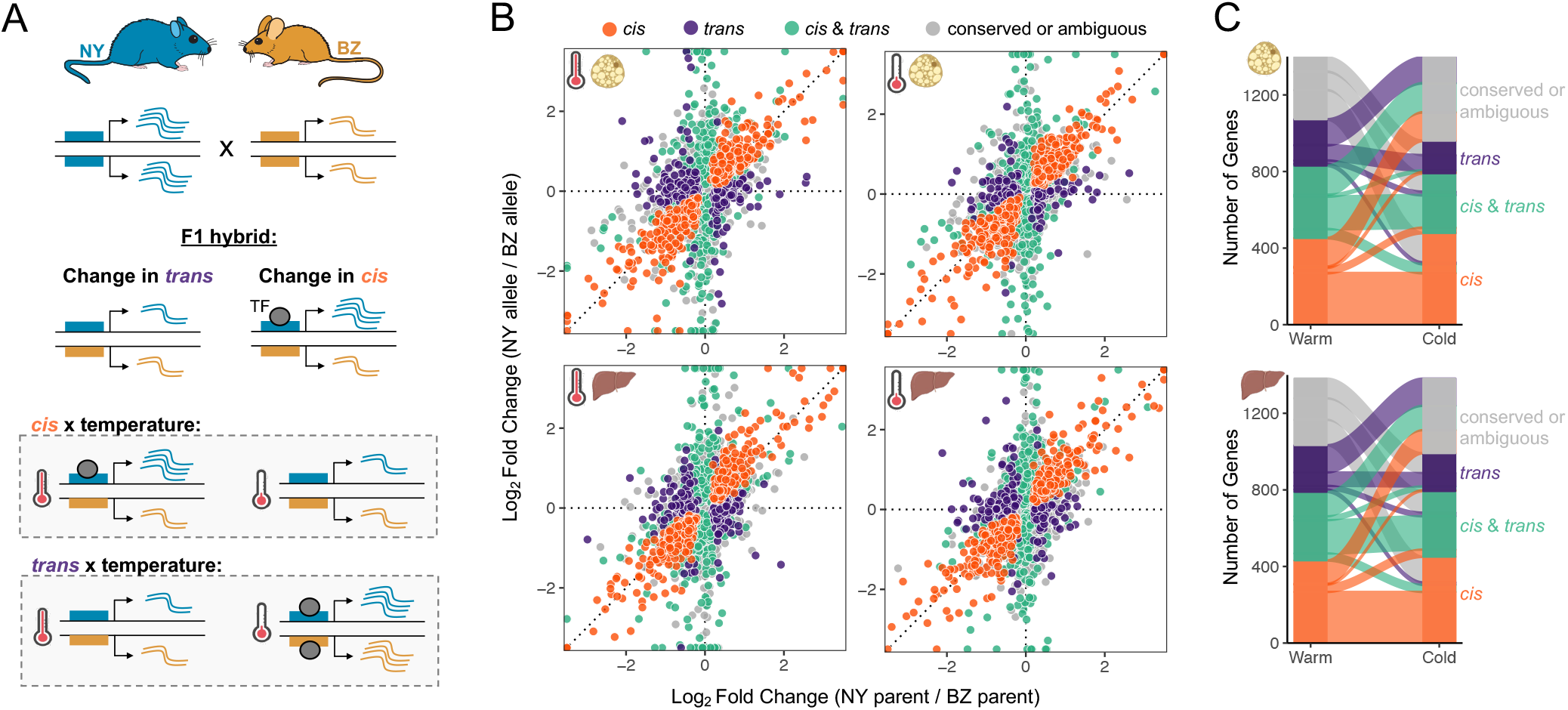
The relative distribution of regulatory changes between New York and Brazil house mice across environments. **(A)** Schematic depicting how *cis*-and *trans*-changes can be inferred with F1 hybrids, and how environmental differences may result in *cis* x temperature and *trans* x temperature effects. Blue and gold boxes represent *cis*-regulatory regions for NY and BZ, respectively. Wavy lines depict transcript levels of an allele. TF = transcription factor. (B) Categorization of regulatory divergence by comparing the expression of NY and BZ parents to NY- and BZ-allele specific expressions within F1s. Points (individual genes) represent log2 fold changes between reads mapping to each allele in the hybrid (BZ allele / NY allele; y-axis) and the reads mapping to each parental line (BZ parent / NY parent; x-axis). Genes are colored based on their inferred regulatory category: orange = *cis*, purple = *trans*, green = *cis&trans*, gray = conserved or ambiguous. Genes categorized as conserved or ambiguous (gray points) constitute roughly 75% of all genes and are centered on the origin and mostly hidden behind other genes. (C) Changes in the number of genes for each inferred regulatory category between temperature regimes are illustrated in the alluvial plot. Genes that were conserved or ambiguous (gray) at both temperatures (∼75%) are depicted in SI *Appendix*, Fig. S7.

We tested 5,898 genes for ASE based on the presence of fixed differences between parental Brazil and New York lines (see Methods). While most genes showed conserved gene regulation between New York and Brazil mice (∼75%), genes with evidence for expression divergence tended to involve changes in *cis* (Fig. 3B). Specifically, 7-8% of genes showed expression divergence due to *cis* alone and 5-6% genes showed evidence of divergence due to *cis* and *trans* (Fig. 3B). Only ∼5% of genes involved regulatory changes solely in *trans* (Fig. 3B). Moreover, the magnitude of *cis*-effects were greater than *trans*-effects per gene (Wilcoxon signed-rank tests; BAT, *P*=2.97 × 10^−27^; liver, *P*=4.64 × 10^−29^). The predominance of *cis*-regulatory changes relative to *trans*-changes is consistent with previous studies in house mice (37–40).

Genes with evidence for *cis*-divergence were enriched for GO terms related to metabolic processes, as well as the pathway for metabolism (Reactome R-MMU-1430728; liver, FDR=6.55 × 10^−8^; BAT, FDR=1.49 × 10^−8^). In the liver, genes with *cis-*regulatory changes showed a greater than 2-fold enrichment of genes with mutant phenotypes for abnormal susceptibility to weight gain (FDR=0.014) and were nominally significantly enriched (with unadjusted *p*-values) for several other phenotypes related to body weight, size, and composition (*SI Appendix*, Fig. S6). Interestingly, two genes (*bcat2, adam17*) exhibiting *cis*-regulatory divergence were previously implicated in body weight differences in North American populations (24), further supporting their role in adaptive divergence between house mouse populations.

### Most *cis*-changes are robust to environmental temperature

We next asked how the environment modulates gene regulatory evolution by comparing patterns of *cis*- and *trans*-regulatory differences across environments. Similar to expression patterns observed in the parents, the majority of genes that could be categorized across temperature treatments showed the same regulatory mode in both environments (∼88%) (*SI Appendix*, Fig. S7). For the genes that did show a change in regulatory mode, we found that *cis*-regulatory changes were more insensitive to temperature than *trans*-changes (Fig. 3C). Comparing the difference in magnitude of the *cis*- and *trans*-differences between warm and cold conditions, we found that *trans*-differences were greater between environments for both tissues (Wilcoxon signed-rank tests, *P* < 2.2 × 10^−16^) (*SI Appendix*, Fig. S8). The cold environment also had a lower proportion of genes with *trans*-divergence (Chi-square tests; BAT, *P*=0.0003; liver, *P*=0.02), where the proportion of genes with only *cis*-divergence was the same across temperature conditions (Chi-square tests; BAT, *P*=0.51; liver, *P*=0.66). These results indicate that *trans*-effects play a larger role in gene expression plasticity than *cis*-effects.

Given that much of gene expression plasticity is governed by changes in *trans*, we next asked whether the observed correlation between plastic and evolved changes (i.e., Fig. 2C) is also seen for genes controlled in *cis*, since expression variation at such genes is not expected to be correlated (41). We therefore compared plastic expression differences with evolved expression differences in genes with evidence for *cis*-regulatory divergence (i.e., ASE) (42). Similar to our previous findings, we found that expression plasticity generally goes in the same direction as evolved divergence in the liver (Spearman’s *ρ* = 0.261, *P* < 0.05) (*SI Appendix*, Fig. S9). However, no correlation was observed in BAT (Spearman’s *ρ* = -0.0374, *P* > 0.05), suggesting that correlated expression patterns in BAT may be regulated by one or a few *trans*-acting modifiers.

### A small number of genes show temperature-dependent *cis*-regulation

While most *cis*-effects were robust to temperature, we were specifically interested in exploring whether any genes showed temperature-dependent *cis*-effects. Such genes are of particular interest since they correspond to *plasticity-*eQTL (i.e., loci that harbor mutations underlying a plastic response)(43). To identify genes for which there was a significant effect of temperature on regulatory divergence, we determined if either the *cis* and/or the *trans* component showed a significant interaction with temperature (see Methods). We identified *cis* x temperature effects for 11 genes in BAT (*gstt1, wars2, hsd11b1, itih5, dst, tmed2, plbd1, cdh13, scd1, tmem45b, s100a13*) and 4 in the liver (*elovl3, hmgcs2, wars2, ebpl*) (FDR < 0.1; 12/15 genes at FDR < 0.05)(*SI Appendix*, Fig. S10). Most of these genes showed differences in the magnitude of ASE between temperatures, but we also observed cases where ASE was induced by one temperature treatment (i.e., *wars2, tmed2, cdh13, s100a13, ebpl, hmgcs2*). Over half of the genes corresponding to *plasticity*-eQTL showed a smaller plastic response in New York than in Brazil, consistent with the overall reduction in expression plasticity in cold-adapted mice. We also identified genes with significant *trans* x temperature effects in BAT (18 genes) and liver (1 gene) (FDR < 0.1; 10/19 genes at FDR < 0.05) (*SI Appendix*, Table S3). Several of these genes with temperature-induced regulatory differences have suggested roles in energy metabolism and thermal tolerance (e.g., refs 44, 45–47).

### Positive selection on genes with *cis*-regulatory divergence in wild house mouse populations

As *cis*-regulatory variants are often drivers of local adaptation (4, 48, 49), and because most regulatory divergence between New York and Brazil house mice is governed in *cis*, we next explored whether genes regulated in *cis* are under positive selection in wild mice from the Americas. To test this, we used previously published whole exome data from wild-caught individuals collected from New Hampshire/Vermont, USA (NH/VT) (21) and Manaus, Brazil (MAN) (Gutiérrez-Guerrero et al., *in prep*), and compared these data to previously published whole genome data from Eurasian populations of house mice (50). Genetic PCA distinguished mice based on subspecies and population-of-origin (Fig. 4A and *SI Appendix*, Fig. S11), with mice from NH/VT clustering most closely with mice from Germany. These results support recent findings that mice from eastern North America are most closely related to populations in northern Europe (51–53).

**Fig. 4.**
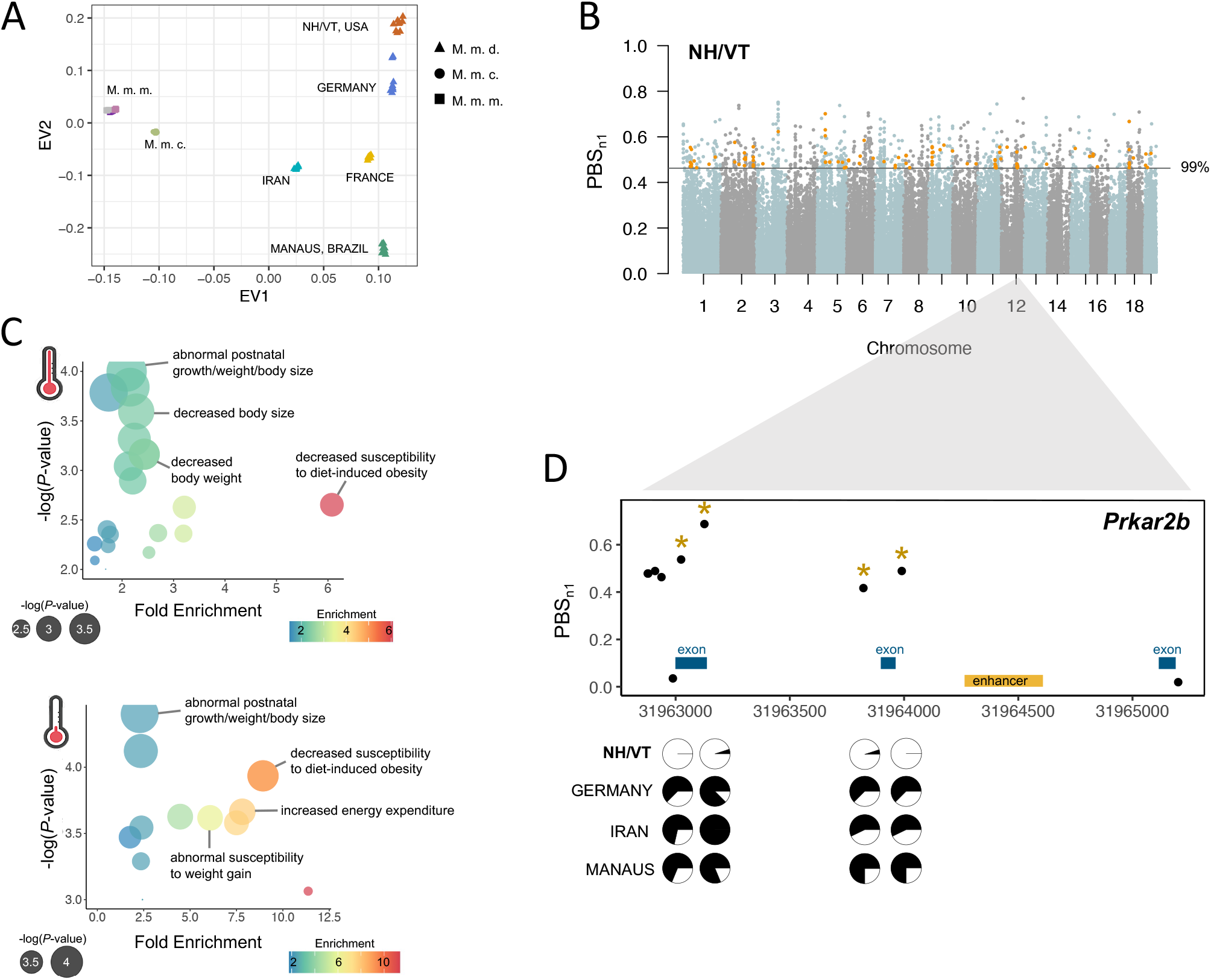
Genomic outliers are enriched in genes with evidence for *cis*-regulatory divergence. **(A)** Genetic PCA of wild house mice distinguished mouse populations based on population-of-origin (*Mus musculus domesticus* (M.m.d.)) and subspecies (*Mus musculus castaneus* (M.m.c.), *Mus musculus musculus* (M.m.m.)). The x and y axes show the first and second SNP eigenvectors, respectively (EV; PC1: 29% of variance, PC2: 8% of variance. **(B)** Autosomal selection scan showing *PBSn1* results for the New Hampshire/Vermont (NH/VT) focal population. Orange points depict genes that exhibit *cis*-regulatory divergence and overlap with outlier regions. **(C)** Gene set enrichment analysis for genes with ASE that overlap genomic outliers in the NH/VT population. ASE outliers were highly enriched for mouse phenotypes related to body size differences and metabolic features, across both temperature treatments. **(D)** Candidate gene that exhibits *cis*-regulatory divergence and overlaps with outlier region. Allele frequencies (pie charts) of significant SNPs (gold asterisks) in the four *Mus musculus domesticus* populations.

Next, to identify genetic signatures of adaptation in house mice from the Americas, we performed a scan for regions of genetic differentiation consistent with selection using a normalized version of the population branch statistic. We used this test to identify highly differentiated loci in our focal populations in the Americas (MAN and NH/VT) relative to Eurasian populations (see Methods). In total, 83,538 and 84,420 non-overlapping 5-SNP windows were analyzed for MAN and NH/VT, respectively. Outlier windows in NH/VT and MAN overlapped 538 and 530 genes, respectively (*SI Appendix*, Dataset S1). Overall, genomic outliers were distributed across the genome in both populations (Fig. 4B and *SI Appendix*, Fig. S12), consistent with selection acting primarily on standing genetic variation in house mice (20, 21).

Finally, we asked to what extent genomic divergence among wild mice from temperate and tropical environments is associated with *cis*-regulatory changes. Specifically, if natural selection associated with climatic adaptation has acted mainly on regulatory variants, we predicted an enrichment of genomic outliers near genes exhibiting ASE (e.g., ref. 54). To test this prediction, we overlapped putative candidate regions for selection based on SNP outlier windows with genes for which we identified evidence for allele-specific expression in BAT or liver. In NH/VT, we found outlier windows overlapped 71 and 62 genes with evidence for *cis*-regulatory divergence under warm and cold conditions, respectively (overlap 44 genes) (Fig. 4B; *SI Appendix*, Dataset S1). The overlap between genes with *cis*-regulatory divergence and outlier windows in this population was greater than expected by chance (hypergeometric test, *P*=0.0016), and therefore is unlikely to be a consequence of genetic drift. Moreover, genes with allele-specific expression were associated with higher average population branch statistic scores than background genes (*P*=0.00026, see Methods). In contrast, we did not find significant overlap between genes with allele-specific expression and genomic outliers for Manaus (*P*=0.4). Outlier windows overlapped 49 and 51 genes with evidence for *cis*-regulatory divergence under warm and cold conditions, respectively (*SI Appendix*, Fig. S12). These genes were not enriched for metabolic process terms or phenotypes.

Genes with allele-specific expression that overlapped genomic outliers in temperate mice were enriched for mutant phenotypes related to body size, growth, and metabolism relative to other genes with *cis*-regulatory divergence (e.g., abnormal postnatal growth/weight/body size, abnormal susceptibility to weight gain, decreased susceptibility to diet-induced obesity, and increased energy expenditure; FDR < 0.05) (Fig. 4C; *SI Appendix*, Dataset S1). This gene set also includes genes whose expression in the liver was previously associated with body mass variation in natural populations of North American house mice (*bcat2, col6a1, col5a2, col3a1*) (21, 24). Additionally, this set included genes implicated in obesity and metabolic phenotypes in humans (e.g., *wrn, plaat3, prkar2b, sulf2, smoc1*) (Fig. 4D) (55) and mice (*SI Appendix*, Table S4). Together, these results suggest that selection has acted on *cis*-regulatory genes related to metabolism and body weight in North American mice (24).

## Discussion

Understanding how both genetic and environmental factors influence gene expression divergence is essential to understanding adaptive evolution. Here, we utilized allele-specific expression in liver and brown adipose tissue to characterize *cis* and *trans* changes underlying expression differences between temperate and tropical house mice when reared under warm and cold laboratory environments. We found that most regulatory divergence was governed by *cis*-regulatory variation, and that these *cis*-effects were largely independent of environmental temperature. However, a subset of genes showed temperature-dependent *cis*-effects and thus represent QTL for expression plasticity. Finally, overlap of genes exhibiting *cis*-regulatory divergence with scans for selection identified several *cis*-regulatory genes under positive selection, consistent with a role for these loci in local adaptation. Together, our results demonstrate how both genetic and environmental effects contribute to adaptive gene expression differences between natural populations.

The rapid colonization of house mice into new environments may have been mediated by plasticity through *trans*-regulation. Specifically, and similar to previous studies, we found that most expression plasticity was largely governed by *trans*-acting factors (43, 56–62), which modify correlated changes in gene expression profiles of hundreds of genes. In fact, a large proportion of the correlated plastic changes we observed went in the same direction as evolved expression divergence, implicating a role for gene expression plasticity in the colonization of new environments (22). Furthermore, this rapid response via *trans*-effects likely shifted to favor the predominant and less pleiotropic *cis*-regulatory architecture over time (62, 63). We found that most *cis*-effects were robust to temperature, indicating a decoupling of environmental plasticity and allelic-effects. Interestingly, a number of these *cis*-regulatory loci show reduced plasticity in temperate mice (*SI Appendix*, Fig. S13), suggesting that selection may target genetic changes that minimize plasticity (30, 32, 64).

Although most *cis*-effects were robust to temperature, we identified a subset of genes that showed temperature-dependent *cis*-effects. These loci are of particular interest since these constitute *plasticity*-eQTL and harbor mutations that directly affect plasticity of gene expression. Genetic assimilation refers to the conversion of a plastic response to a canalized response (65–68). If the ancestral allele at a *plasticity*-eQTL encodes a plastic response and the derived allele encodes a canalized response, then the *plasticity*-eQTL represents a case of genetic assimilation. For example, selection in a cold, temperate environment may have led to the reduced plasticity exhibited in New York mice. A similar mechanism was recently proposed by Verta & Jones (2019) to explain the observed plasticity in gene expression between freshwater and marine threespine stickleback. *Cis*-regulatory variants could rapidly canalize expression through the loss or gain of specific binding sites for conditionally expressed transcription factors, thereby decoupling a gene’s expression from the environment (69). Many of the *cis* x environment candidates illustrate potential regulatory mechanisms underlying genetic assimilation as many of them exhibit reduced plasticity in New York mice (*SI Appendix*, Fig. S13). For example, *scd1* plays an important role in basal and cold-induced thermogenesis (70, 71) and New York mice show higher and constitutive average expression of *scd1* in BAT compared to Brazil mice (*SI Appendix*, Fig. S13). Further study of these genes may help us understand the relationship between plasticity, selection, and adaptation to novel environments in natural populations.

Our comparison between New York and Brazil house mice across environments has implications for our understanding of gene regulation and genome function across short evolutionary timescales. Although house mice colonized the Americas within the last ∼500 years, we found evidence for pervasive *cis-*regulatory divergence. Furthermore, house mice have rapidly adapted to various environments from pre-existing standing genetic variation (20, 21, 24, 72), which has likely contributed to the predominance and enrichment of *cis*-regulatory variation associated with local adaptation in temperate mice. We speculate that the significant overlap between genomic outliers and ASE in temperate mice but not in tropical mice may reflect adaptation predominantly to cold environments (rather than to warm environments), consistent with the warm ancestral range of house mice in the Mediterranean region. Nonetheless, positive selection may preferentially act on *cis*-acting alleles due to their higher additivity and less sensitivity to genomic background (35, 62, 73). Similarly, natural selection may target *cis*-acting alleles due to their insensitivity to environmental conditions, making them a primary substrate for adaptation to novel environments (62). These features of *cis*-regulatory divergence allow them to accrue on extremely short timescales, making them important loci for rapid climatic adaptation.

Overall, this study broadens our understanding of the role of gene regulation in recent adaptive evolution by disentangling *cis*- and *trans*-changes underlying genetic and environmental effects on gene expression differences. While some progress has been made on the relative importance of *cis*- and *trans*-changes in adaptation within and between species (16, 36), most of the observed differences in regulatory patterns have been measured in a single environment, overlooking environment- and genotype-by-environment effects. By pairing allele-specific expression across different conditions with genomic data from natural populations, we discovered important roles for environment-dependent *trans*-changes and environment-independent *cis*-regulatory divergence in populations adapting to new environments. Thus, this work provides insight into the molecular architecture underlying genetic and non-genetic causes of gene expression differences during adaptive evolution.

## Materials and Methods

### Animals and Evolved Phenotypic Differences

To characterize evolved phenotypic differences between New York and Brazil house mice, we used two wild-derived inbred lines of house mice: SARA (New York) and MANA (Brazil). Previous studies have demonstrated that these lines vary in morphology and gene expression and are indicative of population divergence (21, 23) (*SI Appendix, Supplementary Text*). The establishment of these lines has been described previously (23). Mice from each line were housed in a standard laboratory environment at 21^*°*^C with a 12L:12D cycle. Roughly equal numbers of males and females were produced for each within-line comparison (*n* = 32 per line; *SI Appendix*, Dataset S1). We took standard museum measurements on all mice and removed and prepared dried skins. Thermal conductance of pelage (referred to as pelage conductance [W m^-2 *°*^ C^-1^]) was measured on dry skins following the protocol of Riddell et al. 2022 (*SI Appendix, Supplementary Text*) (74). Tail length and ear length were corrected for body mass for each individual. Effects of line and sex for each phenotype were modeled using ANOVA. All statistical analyses were performed using packages available in R (v.4.1.1).

### Experimental Design and Tissue Collection

To investigate the gene regulatory mechanisms underlying local adaptation in house mice, we generated F1 hybrids by crossing a SARA female with a MANA male. All experimental animals were born at room temperature (21^*°*^C) and were provided water and commercial rodent chow *ad libitum*. We weaned and singly housed SARA, MANA, and F1 hybrids at ∼3 weeks of age. We split 3.5-week-old full-sibs and F1 hybrids into size-matched experimental groups across cold (5^*°*^C) and warm (21^*°*^C) treatments. Mice were kept in their respective experimental environment until ∼12 weeks of age, at which point individuals were euthanized via cervical dislocation. We took standard museum measurements and then rapidly dissected and preserved liver and brown adipose tissue in RNAlater at 4^*°*^C overnight and moved to -80^*°*^C until RNA extraction. We prepared standard museum skeletons and accessioned them in UC Berkeley’s Museum of Vertebrate Zoology (catalog numbers are given in *SI Appendix*, Dataset S1). All experimental procedures were in accordance with the UC Berkeley Institutional Animal Care and Use Committee (AUP-2017-08-10248).

### RNA Extraction, Library Preparation, and Sequencing

We extracted total RNA from liver and BAT from each sample (*n* = ∼6 per genotype/sex/treatment/tissue) using the RNeasy PowerLyzer Kit (QIAGEN). We generated Illumina cDNA libraries from 1 *μ*g of purified RNA using KAPA Stranded mRNA-Seq Kit (Illumina), and uniquely indexed libraries using unique dual indexes (Illumina). Libraries were pooled in equal molar concentration and sequenced on one lane each of 150 bp paired-end NovaSeq S1 and NovaSeq S4 at the Vincent J. Coates Genomics Sequencing Center at UC Berkeley. We filtered raw reads below a Phred quality score of 15 and trimmed adapter sequences using *fastp* (75).

### Parental Gene Expression Analyses

After cleaning and trimming parental sequences of MANA and SARA, we mapped reads to the *Mus musculus* reference genome (GRCm38/mm10) using STAR (76). We counted reads overlapping exons using HTSeq (77) based on the Ensembl GRCm38.98 annotation. We imported raw count data into R (v.4.1.1) and transformed expression values using variance stabilizing transformation (78) to assess transcriptome-wide expression patterns via PCA. Next, we removed genes with fewer than an average of 10 reads per individual within each tissue, retaining ∼14K expressed genes per tissue for downstream analyses. We then used DESeq2 (78) on raw, filtered reads to quantify expression patterns by fitting a generalized linear model following a negative binomial distribution (Wald-test). Due to the strong effects of tissue-type and sex on expression patterns (*SI Appendix*, Fig. S1), we computed differential expression between lines for each tissue and sex, separately (but see *SI Appendix, Supplementary Text* for a full parameterized model). Specifically, we used the model line + environment + line*environment to determine the effects of genotype, environment, and genotype-by-environment (GxE) on expression patterns. We defined genes as GxE if: 1) only one genotype showed significant differential expression between temperatures (“line-specific”), or 2) both genotypes showed significant differences between temperatures, but in opposite directions (“opposite”). Lastly, we used a Benjamini-Hochberg multiple test correction (79) on all resulting *P*-values and considered genes with FDR < 0.05 to be significantly differentially expressed.

To determine if gene expression plasticity is correlated with gene expression divergence, we compared genes with significant plasticity in Brazil mice to genes with significant expression divergence between Brazil and New York mice within each tissue and sex, separately. We used Spearman’s rank correlation coefficients to assess overall directionality and significance of gene expression. To account for potential statistical artifacts in the regression (80), we compared the observed correlations to a permuted distribution (10,000 permutations).

### Identifying Variants between Parental Lines

To identify differences between lines for allele-specific read assignment, we performed SNP calling on whole genome sequence data from one female each of MANA and SARA. We mapped genomic reads with Bowtie2 (81) to the mm10 reference genome (setting: –very-sensitive) obtained from Ensembl. We marked duplicates with the Picard tool MarkDuplicates and then we used the GATK tools HaplotypeCaller and GenotypeGVCFs for joint genotyping across genomic samples. We filtered for low quality SNP calls with VariantFiltration (QD < 2.0; QUAL < 30.0; FS > 200; ReadPosRankSum < -20.0). To reduce the influence of genotyping error in whole genome sequencing data on allele-specific expression assignment of RNA-seq reads (e.g., ref. 82), we mapped RNA-seq reads from all individuals and then counted allele-specific reads aligned to each site we genotyped with the GATK tool ASEReadCounter. We excluded sites for which we did not have coverage of at least 5 reads from each population-specific allele. In total, 2,875,480 and 2,181,304 variants from MANA and SARA, respectively, were used for identifying allele-specific reads.

### Mapping Allele-Specific Reads

For allele-specific expression analyses, we mapped reads from hybrid individuals to the mouse reference genome (GRCm38/mm10) using STAR. We used WASP (83) to reduce the potential for reference mapping bias. Specifically, WASP mitigates mapping bias by identifying reads containing SNPs, simulating reads with alternative alleles at that locus, re-mapping these reads to the reference, and then flagging reads that do not map to the same location. Reads that do not map to the same location are then discarded. We retained reads that overlapped a population-specific variant and that passed WASP filtering for our allele-specific expression analysis. We separated reads overlapping informative variants into allele-specific pools (NY, BZ) based on genotype for quantification. We used HTSeq to count the number of reads associated with each gene per population based on the overlap of reads and annotated exonic regions based on the Ensembl GRCm38.98 annotation. We examined per site allelic reads with ASEReadCounter to quantify allele-specific mapping over individual sites. Proportions of reads overlapping the references vs. alternative allele (REF allele / (ALT allele + REF allele)) showed a median 0.5 across samples (*SI Appendix*, Figs. S14-S15), indicating no evidence for reference mapping bias.

### Identifying *Cis- and* Trans-Regulatory Divergence

Parental (F0) and F1 expression data was used to characterize *cis* and *trans* effects. To categorize regulatory divergence at each gene, we inferred differential expression by analyzing raw counts using DESeq2. To identify genes with evidence of allele-specific expression in hybrid individuals, we took reads that mapped preferentially to either New York or Brazil alleles and fit these to a model with allele (NY vs. BZ), sample (individual), and tissue (BAT, liver) for hybrid male samples in DESeq2 (Wald-test; *SI Appendix, Supplementary Text*). As read counts come from the same sequencing library, library size factor normalization was disabled in DESeq2 by setting SizeFactors = 1 for measures of allele-specific expression. We used males to assign regulatory categories to maximize power due to a larger number of hybrid samples sequenced (6 replicates of males vs. 4 replicates of females), though we also see similar regulatory patterns in females (*SI Appendix, Supplementary Text* and Fig. S16). Differential expression between alleles in the F1 is evidence for *cis*-regulatory divergence. Conversely, *trans*-regulatory divergence is inferred when differential expression in the F0 generation is not recapitulated between alleles in the F1. The *trans* component (T) was assessed through a Fisher’s Exact Test on reads mapping to each parental allele in the hybrid vs. parental read counts, summed over all replicates (35, 84). We randomly down-sampled reads to account for library size differences between parental and F1 replicates (39, 85). *P*-values for each test were corrected for FDR with the Benjamini-Hochberg method. We sorted genes into categories based on an FDR threshold of 5% (35, 84), as described below. We analyzed temperature treatments (warm and cold) separately for regulatory assignment and then compared as described below:

*Conserved*: no significant difference between lines (F0), no significant difference between alleles (F1), no significant *T. Cis* only: significant difference between lines (F0), significant difference between alleles (F1), no significant *T*.

*Trans* only: significant difference between lines (F0), no significant difference between alleles (F1), significant *T*.

*Cis & Trans* designations: significant differences between alleles (F1) and significant *T*. This category was further sub-divided into *cis* + *trans* (reinforcing), *cis* + *trans* (opposing), *compensatory*, and *cis* x *trans*, as previously described (37, 39).

#### *Ambiguous*: all other patterns

We identified *cis* x temperature interactions using DESeq2 under a model specifying temperature (cold vs. warm) and allele (BZ vs. NY). To identify *trans* x temperature interactions, we fit a model that included parental and hybrid read counts for temperature (cold vs. warm), allele/genotype (BZ vs. NY), and generation (F1 vs. F0) and interactions. We considered genes significant at an FDR threshold of 10%, consistent with previous studies (e.g., refs. 62, 86) as our statistical analysis has less power to detect interactions than main effects. Similar models were also used to identify sex- and tissue-specific regulatory patterns in DESeq2 (*SI Appendix, Supplementary Text*).

### Genetic PCA of M.m.domesticus populations

We used SNPRelate (87) to perform PCA and IBS hierarchical clustering of population genetic data. Genomic data from 3 Eurasian populations of *M. m. domesticus* (Germany [Cologne-Bonn], France, and Iran) and *M. m. musculus* and *M. m. castaneus* subspecies were downloaded from http://wwwuser.gwdg.de/~evolbio/evolgen/wildmouse/ (50). For PCA, biallelic variants genotyped across all these individuals were extracted and pruned for linkage disequilibrium in SNPRelate (thresholds=0.2) resulting in 22,126 variant sites for PCA and IBS clustering for *M. m. domesticus* comparisons and 25,467 variants for global *Mus* comparisons (Fig. 4A and *SI Appendix*, Fig. S11). Altering the pruning threshold to 0.5 did not result in any change in population clustering.

### Autosomal Scans for Selection

To identify regions with evidence for selection in the Americas, we scanned the exomes of our North and South American focal populations for selection by using a modification of the population branch statistic (PBS) which summarizes a three-way comparison of allele frequencies between a focal group, a closely related population, and an outgroup comparison (*PBSn1*) (88, 89):

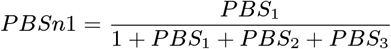

Here, *PBS1* indicates PBS calculated as either Manaus or NH/VT as the focal population, and *PBS2* and *PBS3* indicate PBS calculated for Eurasians populations as the focal populations (France or Germany and Iran, respectively). To maximize the number of sites that could be compared, American populations are not directly compared in the branch test due to the reduced representation of exome data and high per site Fst values between the two populations (*SI Appendix*, Fig. S17). Instead, NH/VT and MAN were each compared to two Eurasian populations [((MAN), France) Iran) and ((NH/VT) Germany) Iran)], selected based on population clustering (*SI Appendix*, Fig. S11). We restricted our SNP set to biallelic variants across the 3 populations being compared and required that at least six individuals in the focal branch be genotyped. We note that the NH/VT sample used in the PBS test is geographically close to the origin of the SARA line.

We used VCFtools (90) to calculate Weir and Cockerham Fst at each variant position. These values were used to calculate *PBSn1* for non-overlapping blocks of 5 SNPs. We consider blocks in the top 1% of *PBSn1* scores outliers and do not attempt to assign *P*-values to each SNP-block (91). Genomic outliers were >3 standard deviations above the mean windowed value of SNP-blocks in each comparison (MAN focal, median=0.045; NH/VT focal median = 0.064). We refer to these genes as “genomic outliers” given the selection scan did not consider neutral models of evolution. We identified windows overlapping genes based on Ensembl gene coordinates (mm10) and the BEDTools “intersect” tool (92). As allele-specific expression in F1s is consistent with local independent genetic changes influencing gene expression, we focused on genes with evidence for *cis*-regulatory divergence (i.e., differences in expression between parental alleles in the F1) for overlap with outlier loci. To ask whether allele-specific expression was associated with elevated *PBSn1* scores, we used a generalized linear model incorporating gene category (ASE or no ASE) and SNP density per kb as factors to *PBSn1* scores. SNP density was calculated by dividing the number of informative sites between NY and BZ for allele-specific expression per gene by transcript length.

## Enrichment Analyses

We performed all GO and pathway enrichment analyses with PANTHER (93, 94). For GO enrichment associated with *cis*-regulatory divergence, we defined the background set of genes as all *cis*-regulated genes tested within a tissue. We performed phenotype enrichment analyses with ModPhea (95), and we annotated genes to specific phenotypes based on Mouse Genome Informatics phenotype annotations (http://www.informatics.jax.org/).

## Data Availability

Scripts are available on GitHub(https://github.com/malballinger/BallingerMack_NYBZase_2023),with the repository archived in Zenodo (accession pending). All sequence data generated in this study have been deposited to the National Center for Biotechnology Information Sequence Read Archive under accession BioProject ID PRJNAXXX. All other data are included in the article and/or *SI Appendix*.

## ACKNOWLEDGMENTS

We thank Beth Dumont for providing whole genome sequences of MANA and SARA. We thank Yocelyn Gutiérrez-Guerrero for providing exome sequences from a population from Manaus, Brazil. We thank Libby Beckman, Eva Fischer, Molly Womack, and four reviewers for helpful feedback on previous versions of the manuscript. Funding and support for this work was provided by the National Institutes of Health (NIH; R01 GM074245 and R01 GM127468 to M.W.N.). This work used the Extreme Science and Engineering Discovery Environment (XSEDE), which is supported by National Science Foundation grant number ACI-1548562. M.A.B. was supported by a National Science Foundation Graduate Research Fellowship (DGE-1106400), a Junea W. Kelly Museum of Vertebrate Zoology Graduate Fellowship, and a University of California Berkeley Philomathia Graduate Fellowship. K.L.M. was supported by a Ruth Kirschstein National Research Service Award from NIH and a Stanford Center for Computational, Evolutionary and Human Genomics postdoctoral fellowship.

## Notes

The authors declare no conflict of interest.

https://github.com/malballinger/BallingerMack_NYBZase_2023

### Competing Interest Statement

The authors have declared no competing interest.

### Summary of Updates

Additional analyses were performed and added to the SI Appendix. Minor textual changes and figure updates were also done with this revision.

