## Supplementary material for "Environmentally robust *cis*-regulatory changes underlie rapid climatic adaptation": SI Appendix

**This PDF file includes:**

Supplementary text
Figures S1 to S19
Tables S1 to S5
Legend for Dataset S1
SI References

**Other supplementary materials for this manuscript include the following:**

Dataset S1

### Supplementary Text

#### 1. Selection of Wild-Derived Inbred Lines

**A. Methods.** For this study, we chose two wild-derived inbred lines (SARA and MANA) as these lines reflect differences that are typical of those seen in other mice sampled from New York and Brazil populations. For instance, gene expression sequenced from both wild-caught and early-generation inbred lines of New York (including SARA, used here) cluster by population of origin, suggesting population divergence (Phifer-Rixey et al. 2018). Moreover, these within-population expression patterns are different from expression patterns seen in lines from other populations. Second, similar patterns are seen at the phenotypic level in both SAR and MAN (Ballinger & Nachman, 2022). For example, multiple lines within SAR (including SARA) are more phenotypically similar to each other than they are to lines in MAN (including MANA, used here). These differences persist across generations (i.e., wild mice to the 10th generation of inbreeding), and are significantly different between SAR and MAN. Furthermore, these phenotypic differences between SAR and MAN are similar to what is seen in wild populations of temperate and tropical house mice, with mice from temperate environments being larger with shorter extremities than mice from tropical environments (Ballinger & Nachman, 2022), suggesting adaptation to different climates. Overall, these results suggest that line-specific differences exhibited by SARA and MANA capture differences seen between populations.

#### 2. Pelage Conductance

**A. Methods.** We measured the thermal conductance of pelage from house mouse specimens (i.e., 'flat skins'; $n$ = 32 per line) to determine differences between populations from New York and Brazil. We sacrificed experimental mice and then air dried the specimens for at least two weeks until completely dry. We used a device that combines a heat flux transducer with controlled heat dissipation across flat skin specimens to measure the thermal conductance of the pelage (see Riddell et al. 2022 for a detailed explanation of methods).

Briefly, the device consisted of a copper pipe (2.54 cm in diameter) that circulated warm water controlled at 37°C to simulate the normothermic body temperature of small mammals. The water was circulated through the pipe using a submersible water pump (Aquatop, N-302) from a temperature-controlled bath into the copper pipe. The copper pipe was embedded into a 6-inch cube of polystyrene to minimize heat transfer away from the pipe, and the tip of the copper pipe was flush with the foam insulation to direct the flow of heat from the warm copper pipe into the flat mammal skin positioned on top of the copper pipe. Between the copper pipe and specimen, we placed a heat flux sensor (FluxTeq, PFHS-01) on top of the copper pipe and secured using double-sided tape. The heat flux sensor (surface area = 1.6 $cm^2$) was a differential-temperature thermophile made of Kapton (polyimide) with a sensor thickness of approximately 305 microns. The heat flux sensor contains a type-T thermocouple embedded into the sensor to simultaneously measure heat flux (W $m^{-2}$) and temperature (°C) across the sensor surface. Then, we applied a thin layer of petroleum jelly (Vaseline™) to the skin on the underside of the specimen to ensure that the skin was sealed to the heat flux sensor without any air pockets (Walsberg et al. 1978). We then clamped the edges of the specimen to further ensure a lack of air pockets using DeWalt® Trigger Clamps while also not disrupting the section of pelage being measured. At the tips of the pelage, we placed a precision wire type-T probe thermocouple (Thermoworks, PT-6; gauge = 0.07366 cm) using an alligator clip to measure the temperature gradient across the mammal pelage. The entire apparatus was placed inside a temperature-controlled incubator chamber (ReptiPro 6000) held at 30ºC.

We recorded the heat flux and temperature from various thermocouples using an analog voltage measurement system (FluxTeq, FluxDAQ) with an integrated thermistor for cold junction temperature compensation. The FluxDAQ continuously measured the temperature of the copper pipe, heat flux across the copper pipe into the skin, temperature above the pelage, air temperature, and wall temperature of the incubator. We used air temperature and wall temperature measurements to ensure black-body conditions within the chamber (1). We measured a single estimate of conductance across the dorsal midline of each specimen, with each measurement taking roughly 30 – 45 m to come to thermal equilibrium. After the readings had come to equilibrium, we recorded the average conductance value over 90 s. We then calculated thermal conductance (W $m^{-2}$ $°C^{-1}$) by dividing the heat flux from the heat flux transducer (W $m^{-2}$) by the difference in temperature (°C) between the heat flux transducer and tip of the pelage (Walsberg 1988).

#### 3. Extended Parental Gene Expression Analyses

**A. Methods.** To determine the effects of sex on gene expression evolution, we quantified genotype-by-sex interactions (GxS) by fitting a generalized linear model following a negative binomial distribution using DESeq2 (Wald test; Love et al. 2014). Specifically, we determined the effects of genotype, sex, and genotype-by-sex on expression patterns for each tissue and temperature treatment, separately. We removed genes with fewer than an average of 10 reads per individual within each tissue. We used a Benjamini-Hochberg multiple test correction (Benjamini and Hochberg, 1995) on the resulting *P*-values and considered genes with a false discovery rate (FDR) smaller than 0.05 to be significantly differentially expressed.

To determine the effect of tissue, sex, genotype, environment, and their interactions on parental gene expression, we fit a fully parameterized linear model using DESeq2 (~ tissue + sex + line + environment + tissue:sex + tissue:line + tissue:environment + sex:line + sex:environment

+ line:environment + tissue:sex:line + tissue:sex:environment + sex:line:environment + tissue:sex:line:environment). We removed genes with fewer than an average of 10 reads per individual. We used a Benjamini-Hochberg multiple test correction on the resulting *P*-values and considered genes with an FDR smaller than 0.05 to be significantly differentially expressed. We also compared this full model to a reduced model without the four-way interaction using a likelihood ratio test to directly determine the effect of tissue-by-sex-by-genotype-by-environment interactions (TxSxGxE) on parental gene expression (Love et al. 2014).

**B. Results.** Similar patterns in expression divergence and expression plasticity were seen across both males and females (Figures 1E and S2-S3). However, notable differences in plasticity were observed in females, especially in BAT. First, tissue-specific differences in plasticity were evident through hierarchical clustering, as female Brazil samples clustered more strongly by genotype and environment in liver but not in BAT (Figure S18). Second, these clustering differences are largely driven by plastic responses in Brazil females. For instance, roughly 40 genes showed opposite divergence patterns in cold versus warm environments (Figure S3A), with Brazil females exhibiting higher expression in the cold. Furthermore, most of these genes exhibit GxE (Figure S3B), with patterns of expression plasticity aligning with evolved divergence (Figure S3C). Interestingly, the majority of these genes are myosin-related or associated with muscle function which has broader implications for at least two reasons. First, BAT is developmentally and functionally more closely aligned to skeletal muscle than it is to white adipose tissue (Timmons et al. 2007; Forner et al. 2009). Second, myosin-regulated genes recruited by BAT have been shown to promote and enhance nonshivering thermogenesis under cold conditions (Liu et al. 2023; Tharp et al. 2018). Thus, these expression patterns in cold Brazil females suggest increased thermogenic activity.

Although males and females showed similar patterns of gene expression evolution, we asked whether differences in sexual dimorphism in gene expression also played a role in expression divergence between New York and Brazil. To characterize the role of genotype-by-sex interactions in expression divergence, we identified GxS for each tissue and environment separately (see above Methods). Consistent with strong patterns of divergence, the vast majority of genes showed differential expression between New York and Brazil mice, regardless of sex (Figure S4). Interestingly, however, BAT harbored very little GxS compared to liver (Figure S4). When we examined sample-to-sample distances with hierarchical clustering, BAT samples indeed clustered strongly by genotype but not by sex (Figure S18).

Finally, we determined the effects of tissue-type, sex, genotype, environment, and their interactions on parental gene expression using a full parameterized model (Table S5). Tissue-type showed the greatest number of differentially expressed genes (13,713) followed by genotype (8,480; Table S5). We did not detect any significant TxSxGxE effects on parental gene expression ($P > 0.05$).

#### 4. Extended Gene Regulatory Divergence Analyses

##### A. Methods.

***Regulatory Categorizations in Females.*** To determine if patterns of *cis*- and *trans*-regulatory divergence are similar in both males and females, we examined regulatory divergence in females subjected to cold-treatment to maximize power due to a larger number of hybrid samples sequenced (6 replicates of cold females vs. 4 replicates of warm females). We categorized *cis*- versus *trans*-regulatory divergence at each gene using DESeq2 (see main manuscript Methods for details).

***Sex-Specific Gene Regulatory Patterns.*** To understand the relative contribution of sex differences to *cis*- and *trans*-regulatory divergence between New York and Brazil, we assessed the extent to which regulatory control is sex-biased. Males were used to assign regulatory categories to maximize power due to a larger number of hybrid samples sequenced (6 replicates of F1 males vs. 4 replicates of F1 females). Sex-specific allele-specific expression was identified for autosomal genes in DESeq2 under the model ~sex + individual:sex + allele:sex by contrasting male:allele and female:allele based on a randomly chosen equal number of male and female F1s. To identify *trans**sex interactions, we fit a model that included parental and hybrid read counts (~allele + allele:generation + population:sex + population:generation:sex) for each tissue separately for autosomal genes.

We note that interactions between X-linked *trans*-factors and autosomal *cis*-regulatory variants, which have been documented in crosses between inbred lines of *Drosophila melanogaster* (e.g., Coolon et al. 2013), can contribute to variation in gene expression. As males used in this study all have the same X chromosome (SARA), we were unable to identify regulatory divergence in males due to interactions between the MANA X and *cis*-regulatory polymorphisms.

***Tissue-Specific Gene Regulatory Patterns.*** While both liver and BAT play essential roles in metabolism and thermogenesis, these tissues have distinct functional properties that differentiate their role in environmental adaptation. Thus, we assessed the extent to which regulatory control is tissue-biased. Similar to sex-specific gene regulatory patterns described above, we used males to assign regulatory categories to maximize power due to a larger number of hybrid samples sequenced. Tissue-specific allele-specific expression was identified for autosomal genes in DESeq2 under the model ~tissue + individual:tissue + allele:tissue by contrasting liver:allele and BAT:allele for each temperature separately.

##### B. Results.

***Female-Specific Regulatory Divergence.*** Similar to gene regulatory patterns identified in males (see Figure 3B), most genes showed conserved gene regulation between New York and Brazil females (~79%). As in cold-treatment males, the majority of regulatory divergence was found to be a result of *cis*-regulatory changes (Figure S16). *Cis*-divergence alone accounted for 8.4% and 7.9% of genes with regulatory divergence in BAT and liver, respectively, whereas *trans*-only changes accounted for 4.2% and 4.4% of genes. Comparing regulatory designations across males and females for individual genes, the majority of genes showed conserved divergence across the sexes (~84% for BAT and for liver under cold conditions). In the majority of cases where genes were found in different regulatory categories between sexes, genes were found to have "conserved" or "ambiguous" regulation in one sex but not the other (~74%), which may in part due to differences in power in addition to sex-biased differences in regulation (see "Sex-Specific Regulatory Divergence" results below).

***Sex-Specific Regulatory Divergence***. To understand the relative contribution of sex differences to regulatory divergence, we compared the differences in effect sizes between males and females (i.e., average | male effect size *cis* - female effect *cis* | vs. average |male effect size *tran*s - female effect size *trans* |). Sex was found to have a larger average effect on *trans* divergence in both tissues ($P < 2.2 \times 10^{-16}$). Next, we investigated genes for sex-specific allele-specific expression. We found limited evidence for sexually dimorphic gene regulatory divergence between lines. Contrasting allelic expression between males and females, we identified 22 autosomal genes in the liver and 11 genes in BAT with sexually dimorphic allele-specific expression (FDR < 0.1). Six of these genes showed sexually dimorphic allele-specific expression in both tissues (*ebpl, ddx55, fh1, tmed2, spata13, C130074G19Rik*). Comparing expression between male and female individuals of the parental and hybrid generation, we also identified 26 autosomal genes with a significant effect of sex on *trans*-divergence (FDR < 0.1).

***Tissue-Specific Regulatory Divergence.*** Comparing gene expression evolution in BAT and liver, we found regulatory divergence to be largely tissue-biased. The majority of genes (80%) for which we could assign a regulatory category in each tissue were assigned to a different regulatory category in the other tissue (2954/3672 genes). In particular, we found that *trans*-divergence was more likely to be restricted to one tissue (with expression conserved between lines in the other tissue), compared to *cis*-changes which were more often shared (>2-fold more) (Chi-square test $P < 0.0001$). This may reflect the general observation of increased tissue-specificity of *trans*-effects relative to *cis*-effects (GTEx Consortium 2017).

To formally identify tissue-biased ASE, we contrasted ASE measurements in BAT and liver for paired hybrid samples (i.e., differential allele-specific expression). We identified 338 genes with evidence for differential allele-specific expression between tissues (Figure S19). *Cis*-effects are not independent across tissues, as ratios of BZ to NY read counts are correlated between tissues, suggesting shared regulatory divergence between BAT and liver (Spearman's rho = 0.44; Figure S19). Furthermore, while the majority of these genes (77%) showed significant allele-specific expression in just one tissue, we also identified cases where allele-specific expression was present in both tissues but with differences in expression magnitude or direction (23%). Of these genes, forty-three had discordant allele-specific expression between tissues, where the opposite parental allele was up-regulated between tissues. Genes with tissue-biased ASE were enriched for metabolic phenotypes (e.g., abnormal lipid homeostasis, FDR=0.00027; increased food intake, FDR=0.036) and tissue specific functions and physiology (e.g., abnormal adipose tissue physiology, FDR=0.007; abnormal liver morphology, FDR=0.00097). These results highlight the importance of tissue-specific gene regulation in population divergence (e.g., Hart et al. 2018; Verta & Jones, 2019).

#### 5. Comparison of DESeq2 to Traditional Binomial Tests

**A. Methods.** To compare our method of categorizing genes with *cis-* vs. *trans*-divergence utilizing DESeq2 to the traditional approach of binomial exact tests (e.g., McManus et al. 2010, Coolon et al. 2014, Lemmon et al. 2014, Mack et al. 2016, Hu et al. 2022), we followed the approach of Coolon et al. (2014) and Mack et al. (2016). Briefly, using reads from BAT under warm conditions, we down sampled to equalized power between F1 and parental samples and pooled across replicates. Binomial exact tests using the R base function binom.test were then used to identify allele-specific expression (F1: NY vs. BZ allele) and differential expression between parents (F0: NY parent vs. BZ parent) and divided genes into regulatory categories based on FDR thresholds as described in the main text (*cis*-only, *trans*-only, *cis*+*trans* divergence, and conserved/ambiguous expression patterns).

**B. Results.** Comparing across tests, the majority of genes fell within the same regulatory category (91.4%; 5,384 genes). The predominance of *cis*-regulatory changes (7.7% and 9% of genes with DESeq2 and binomial approach, respectively) over *trans*-changes (4% and 3.5% of genes with DESeq2 and binomial approach, respectively) was consistent between methods. Binomial and DESeq2 *p*-values were found to be highly correlated (Spearman's rank correlation, *rho*=0.93, $P < 2.2\times10^{-16}$).

Of the genes that were categorized differently between approaches, the majority were categorized as conserved/ambiguous under one method and divergent under the other (87% of discordant genes). In these cases, binomial tests more often categorized genes as divergent where the utilization of DESeq2 resulted in a gene being identified genes as conserved (74%). Comparing genes with non-conserved regulatory assignments in at least one analysis, we did not observe a significant relationship between read depth and discordant category assignment between approaches overall (Wilcoxon Test, *P*=0.75).

<u>**Supplemental Figures:**</u>

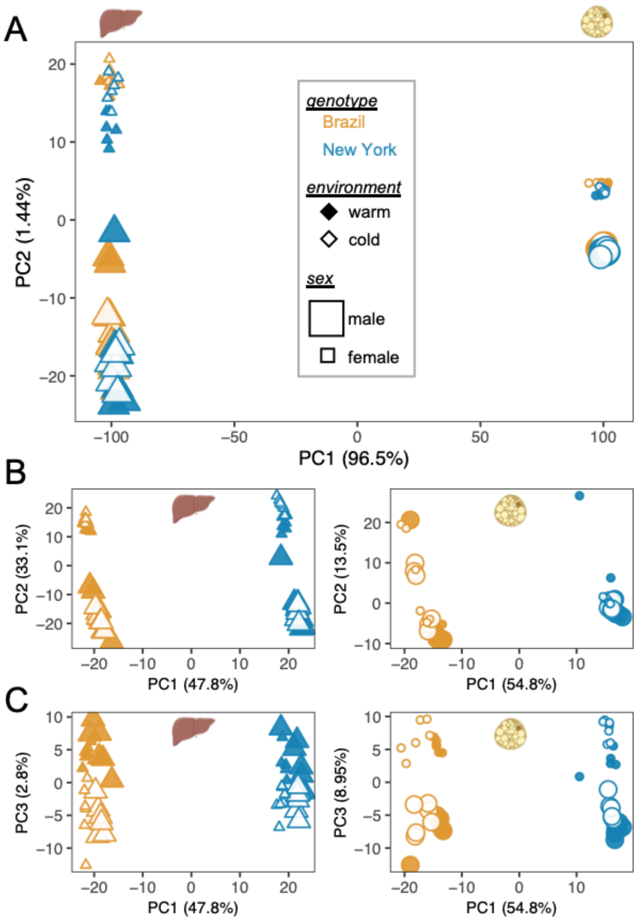


<u>**Figure S1:**</u> **Principal components analysis explaining expression level variation of Brazil and New York mice across tissue, sex, and environment. (A)** When analyzing all expression data together, PC1 explains ~97% of the variance and reflects tissue-type, while PC2 explains ~1.5% of the variance and reflects differences in sex. **(B)** When analyzing expression data for each tissue separately, PC1 explains ~50% of the variance and reflects genotype differences for both tissues. While PC2 reflects sex differences in the liver, there are no clear patterns associated with PC2 in BAT. **(C)** PC3 explains ~3% of the variance in liver and reflects differences in environment. In BAT, PC3 explains ~9% of the variance and reflects differences in sex. Overall, within each line, sex caused substantial differences in gene expression, but along different PC axes with each tissue.

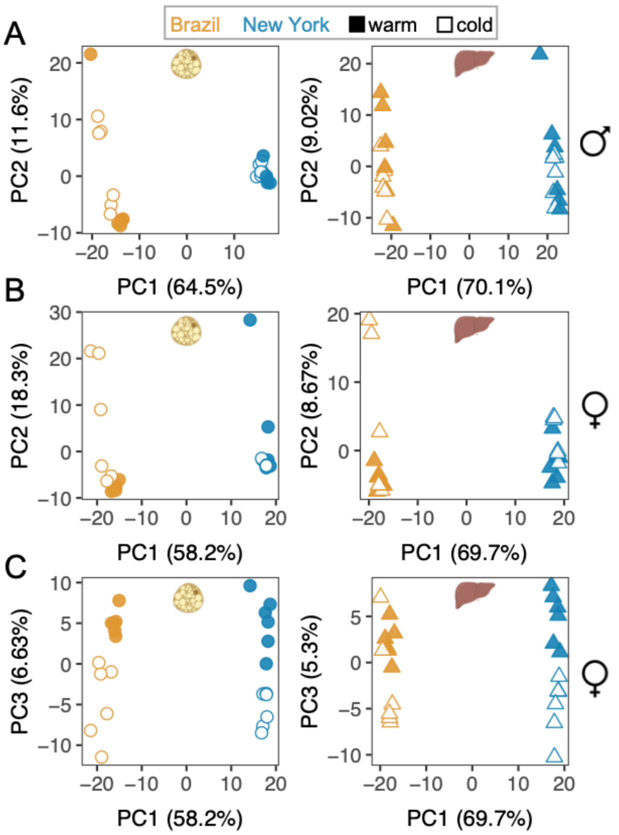


**<u>Figure S2:</u> Principal components analysis explaining expression level variation of Brazil and New York mice across environments for tissue-type and sex, separately.** While PC1 explains roughly 60% of the variance in both male **(A)** and female **(B)** gene expression (and reflects genotype differences), PC2 does not clearly separate individuals by environment. **(C)** In females, PC3 explains > 5% of the variance in both tissues, and largely separates out samples based on environment across both tissues.

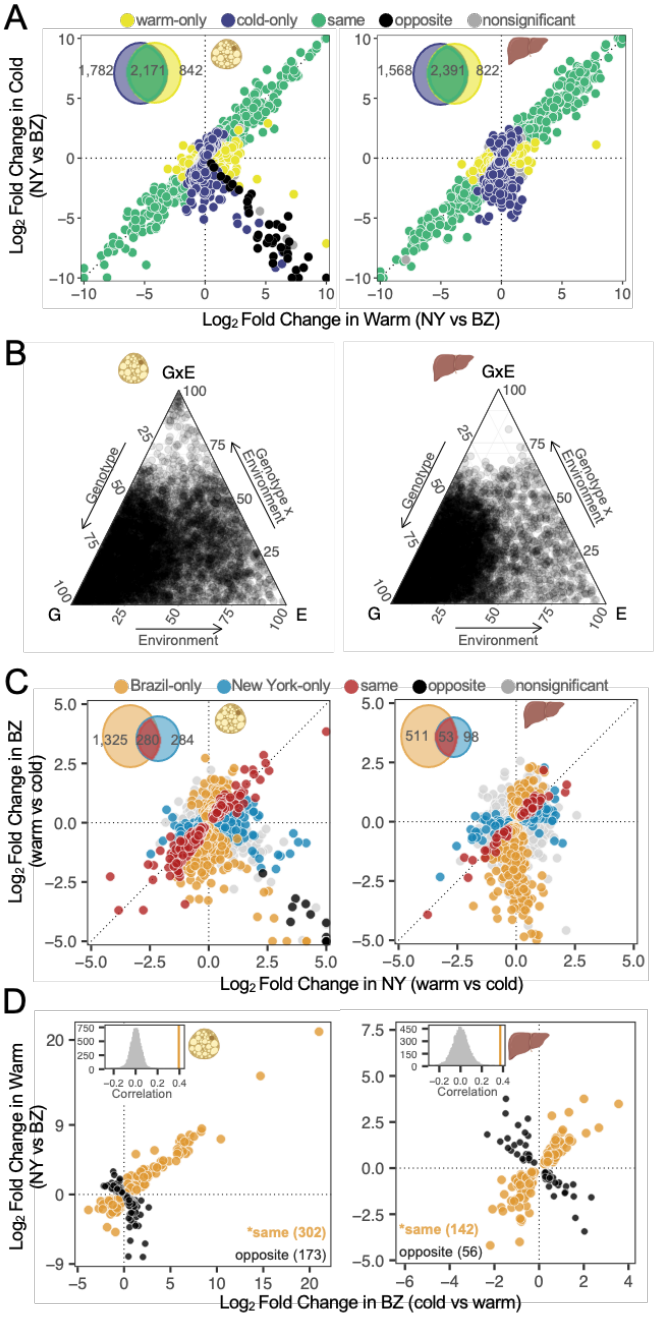


**<u>Figure S3:</u> Divergence and GxE patterns in females.** **(A)** Expression divergence between temperature-regimes in BAT and liver. Log2 fold changes between parents were calculated for all genes independently. In each panel, genes (points) are colored depending on their direction and significance of the log2 fold change (FDR < 0.05). Thirty-nine genes show opposite patterns in BAT, with higher expression in both New York warm and Brazil cold. **(B)** Ternary plots depicting the proportion of each gene's expression variance explained by genotype (G), environment (E), and GxE. The relative proportion of each factor is shown for all differentially expressed male genes in BAT and liver. Total variance is the sum of all three components. **(C)**

Comparison of gene expression differences between temperature regimes in NY and BZ females in both BAT and liver. Log2 fold changes between temperatures were calculated for all genes independently. In each panel, genes (points) are colored depending on their direction and significance of the log2 fold change. GxE categories include line-specific responses or opposite responses between lines. Insets depict the total number of differentially expressed genes for each comparison (FDR < 0.05). **(D)** The relationship between plasticity of gene expression and evolved divergence in gene expression in BAT and liver. Each gene (point) represents expression differences with statistically significant plasticity in Brazil (cold vs warm; FDR < 0.05) as well as significant expression divergence between NY and BZ at warm temperature (FDR < 0.05). Points colored in orange represent genes with a positive correlation between plasticity and evolved divergence. Points in black represent genes with a negative correlation. Insets depict the observed correlation coefficient (orange solid lines) are more positive than a randomized distribution of correlation coefficients for each tissue (see main manuscript Methods for details). Asterisks denote significance of adaptive plasticity for each tissue (binomial exact tests, $P < 0.05$).

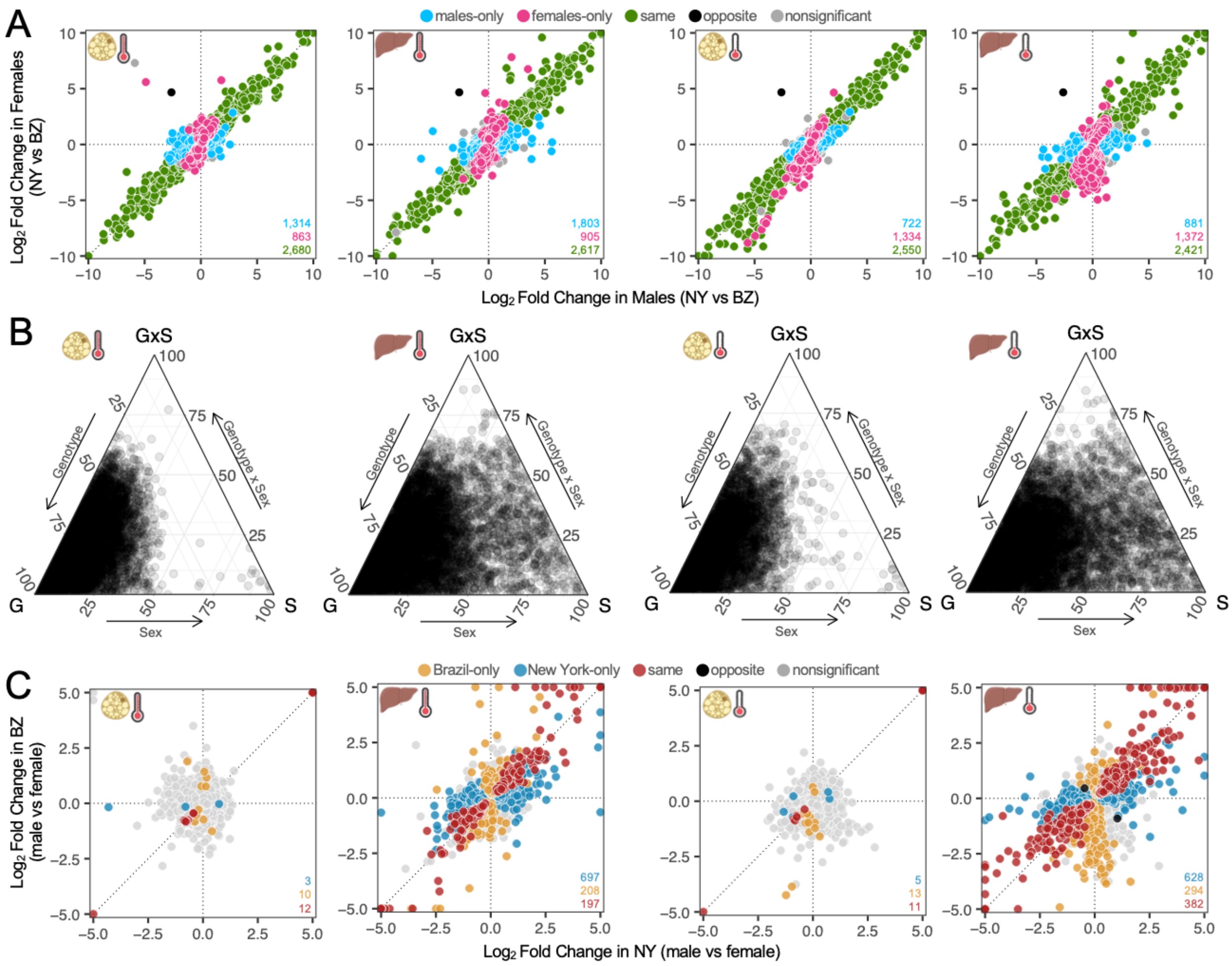

**<u>Figure S4:</u> Parental expression patterns of divergence and genotype-by-sex interactions (GxS). (A)** Expression divergence between New York and Brazil across males and females (both tissues and temperature treatments). Log2 fold changes between parents were calculated for all genes independently. In each panel, points (representing individual genes) are colored depending on their direction and significance of the log2 fold change (FDR < 0.05). Insetted numbers portray the total number of differentially expressed genes for each comparison (FDR < 0.05). **(B)** Ternary plots depicting the proportion of each gene's expression variance explained by genotype (G), sex (S), and GxS. The relative proportion of each factor is shown for all differentially expressed genes in BAT and liver across both temperature treatments. Total variance is the sum of all three components. **(C)** Comparison of gene expression differences between sexes in NY and BZ mice in both BAT and liver and across both temperature treatments. Log2 fold changes between sexes were calculated for all genes independently. In each panel, points are colored depending on their direction and significance of the log2 fold change. GxS categories include line-specific responses or opposite responses between lines. Insetted numbers portray the total number of differentially expressed genes for each comparison (FDR < 0.05).

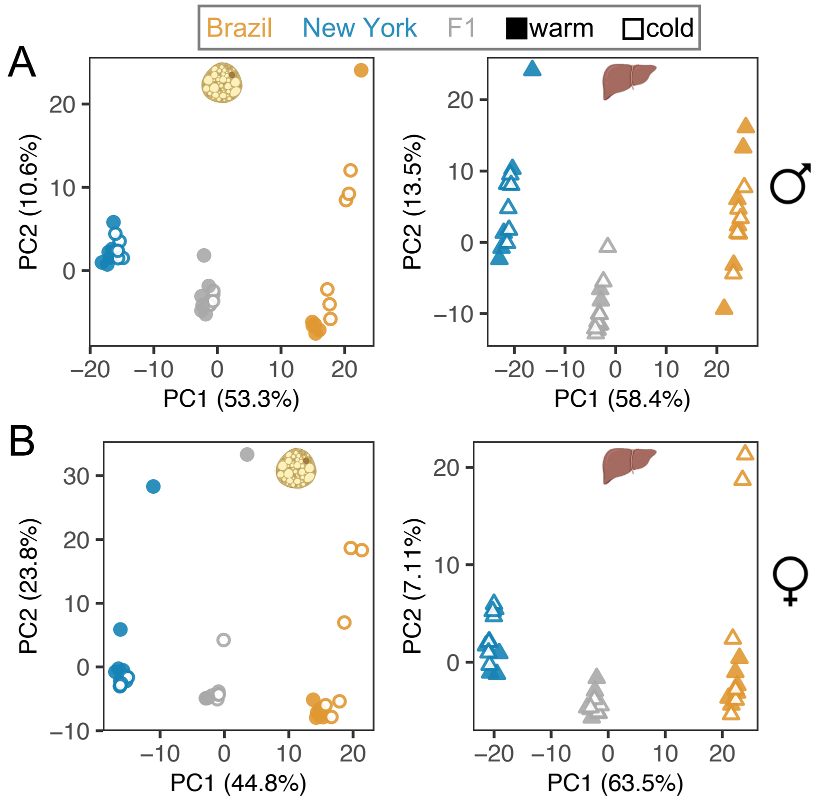


**<u>Figure S5:</u> Principal components analysis explaining expression level variation of Brazil, New York, and F1 hybrid mice for each tissue and sex separately.** In males (A) and females (B), PC1 explains > 40% of the variance in both tissues and cleanly separates out individuals by genotype.

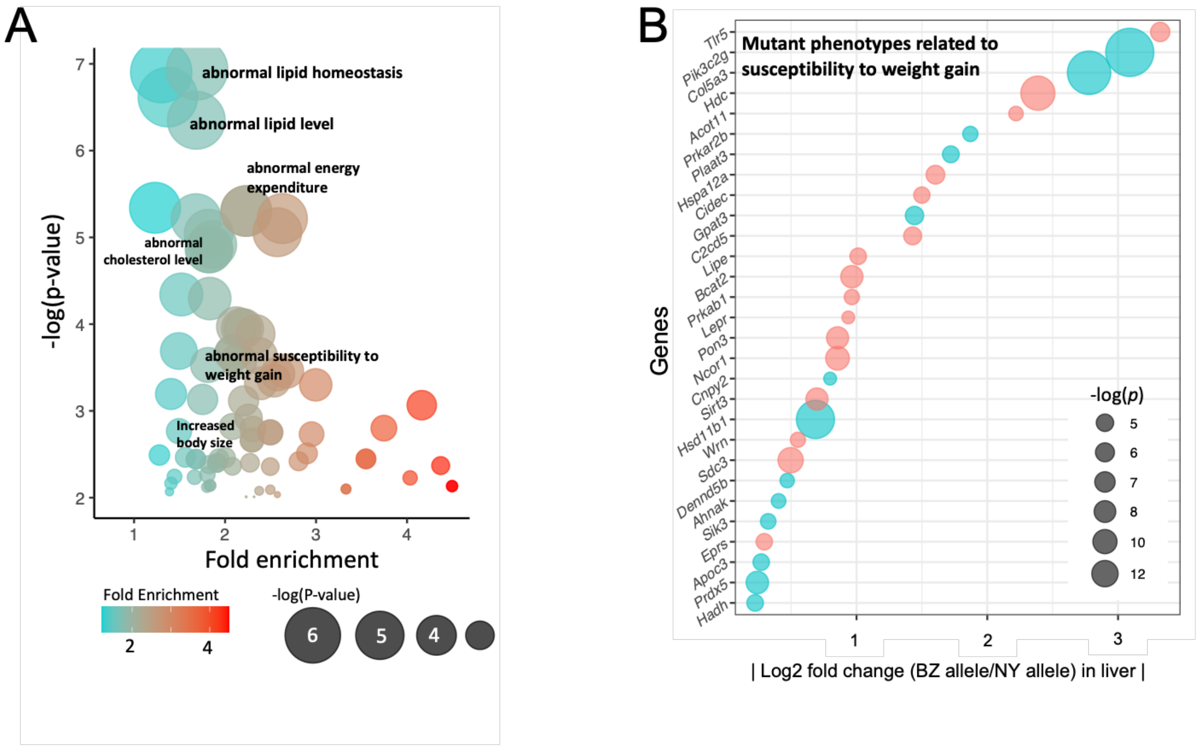


**<u>Figure S6:</u> Enrichment of *cis*-regulatory changes in the liver. (A)** Enrichment plot depicting liver genes with *cis*-regulatory changes for various mutant phenotype annotations for homeostasis and metabolism. Size of circle denotes significance while color denotes fold enrichment. **(B)** Candidate liver genes that show *cis*-changes and have a 2-fold enrichment with mutant phenotypes related to susceptibility to weight gain. Size of circle denotes significance while color of circle denotes higher expression of either BZ allele (blue) or NY allele (red).

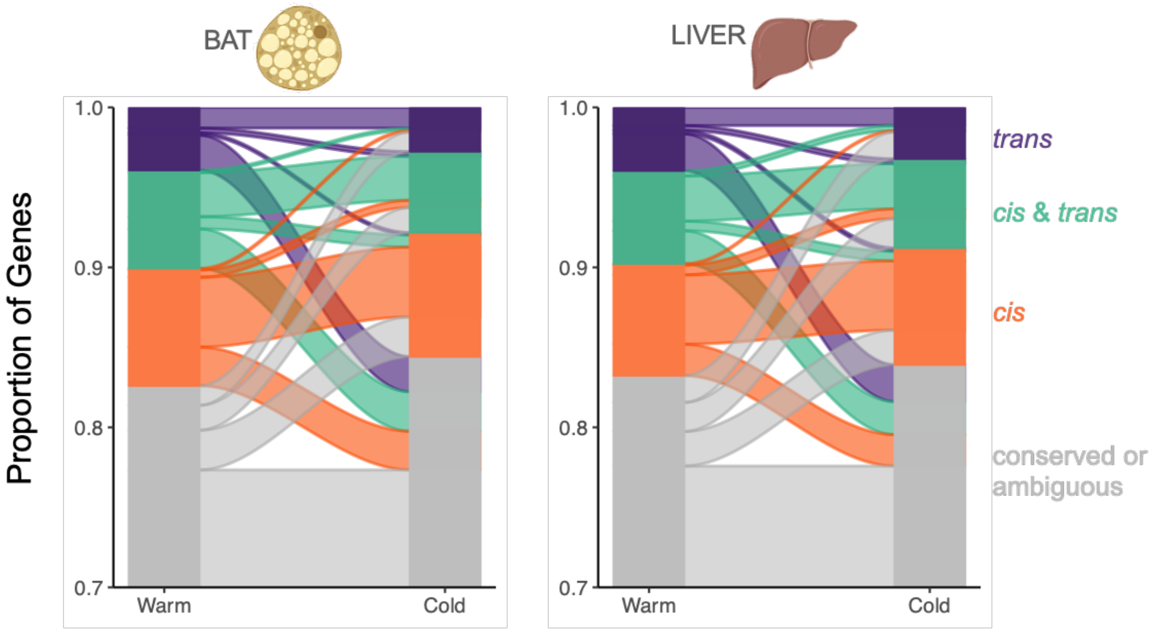


**<u>Figure S7</u>. Gene regulatory changes between New York and Brazil house mice across environments.** Changes in the number of genes for each inferred regulatory category between temperature regimes are illustrated in the alluvial plot. Note: given that ~75% of genes are conserved+ambiguous and do not change between environments, the y-axis begins at 70%.

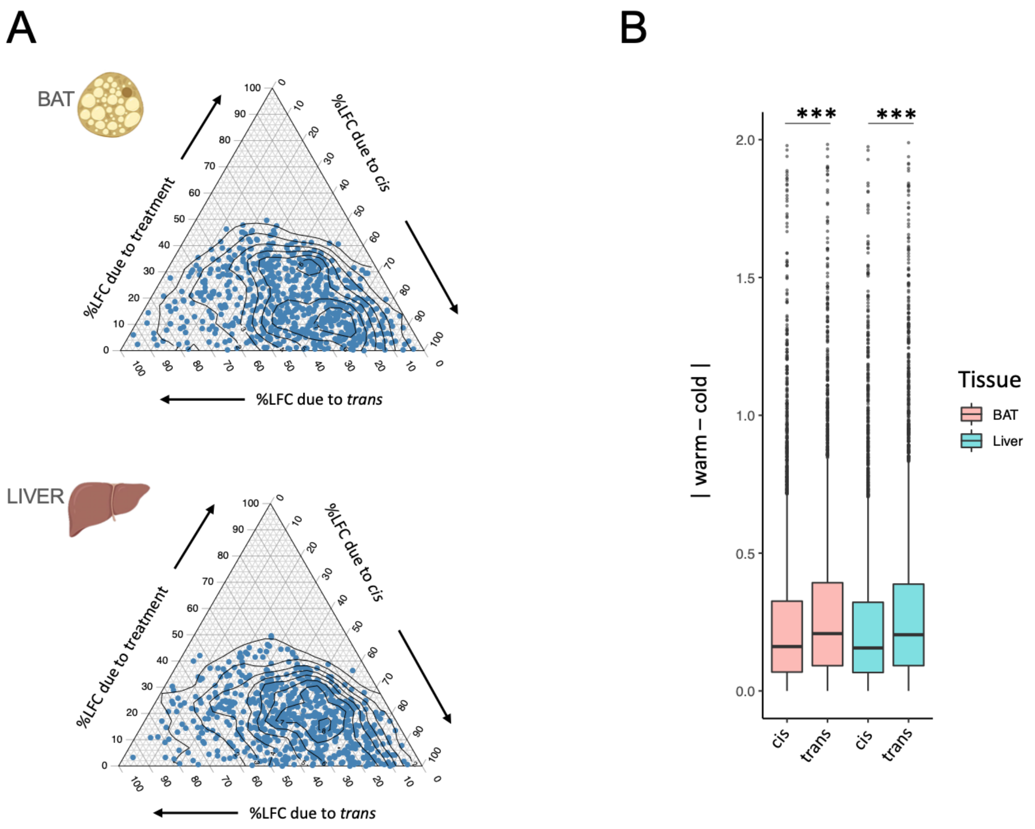


**<u>Figure S8.</u> Effect sizes of *cis*- and *trans*-changes between environments for both BAT and liver. (A)** The relative proportions of log2-fold change due to *cis*, *trans*, and temperature for genes with significant genetic effects (*cis* and/or *trans*) in BAT and liver (see Methods for statistical identification of *cis*- and *trans*-effects). Total variance is the sum of all three components. **(B)** Boxplots displaying absolute log2-fold change of *cis*- and *trans*-difference between warm and cold environments, with *trans*-difference being greater across environments for both tissues (Wilcoxon signed-rank tests, ***$P < 0.001$).

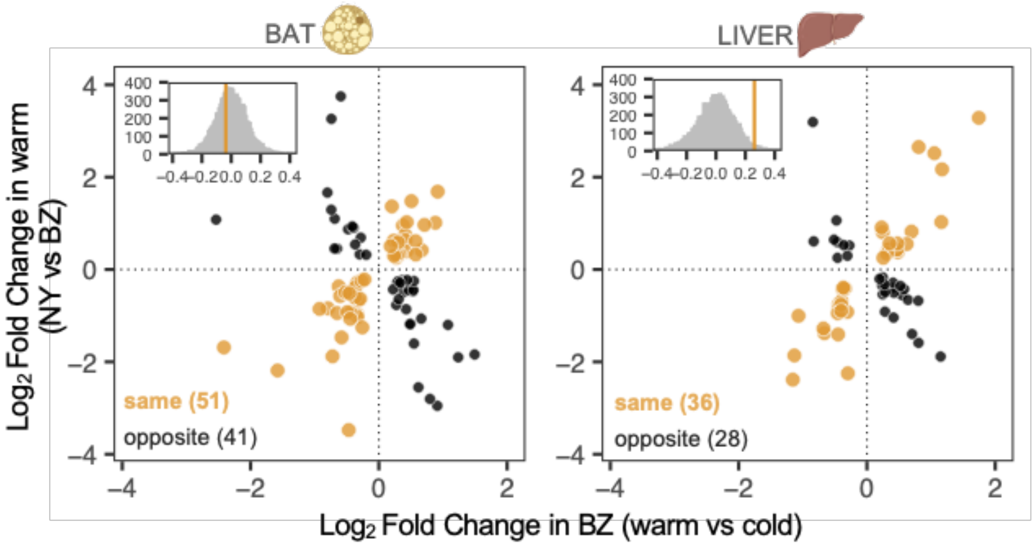


**Figure S9. The relationship between plasticity and evolved divergence in *cis*-regulated genes.** Each *cis*-regulated gene (point) represents expression differences with statistically significant plasticity in Brazil (cold vs warm; FDR < 0.05) as well as significant expression divergence between NY and BZ at warm temperature (FDR < 0.05). Points colored in orange represent genes with a positive correlation between plasticity and evolved divergence. Points in black represent genes with a negative correlation. Insets depict the observed correlation coefficient (orange solid lines) on a randomized null distribution of correlation coefficients for each tissue (see main manuscript Methods for details).

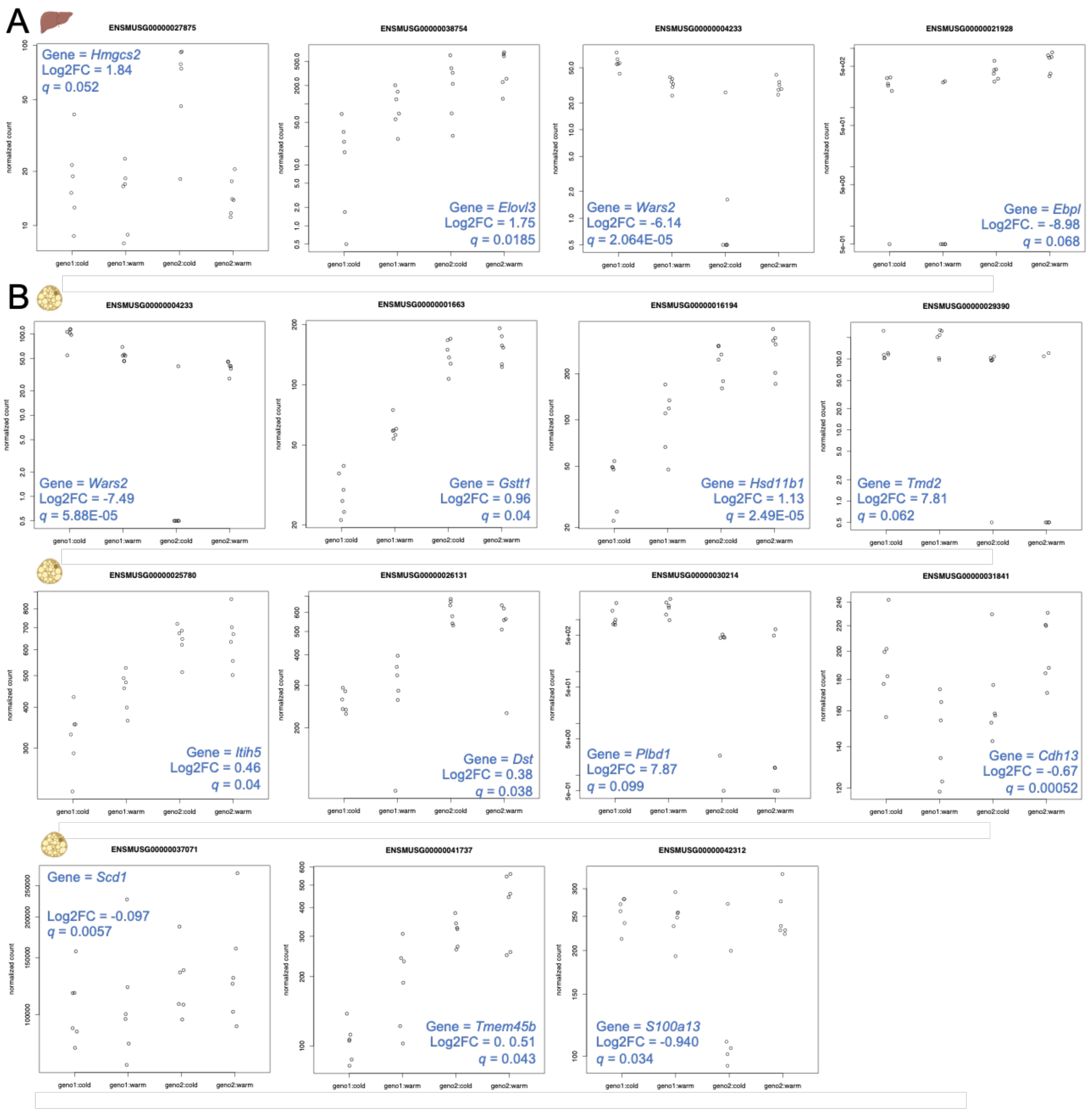


**<u>Figure S10.</u> Effect sizes for the 15 genes showing significant *cis* x temperature effects.** Each plot (which represents a single gene) includes F1 counts, associated *p*-values, and log2-fold changes for interactions. Top row depicts 4 liver genes, while bottom three rows depict 11 BAT genes.

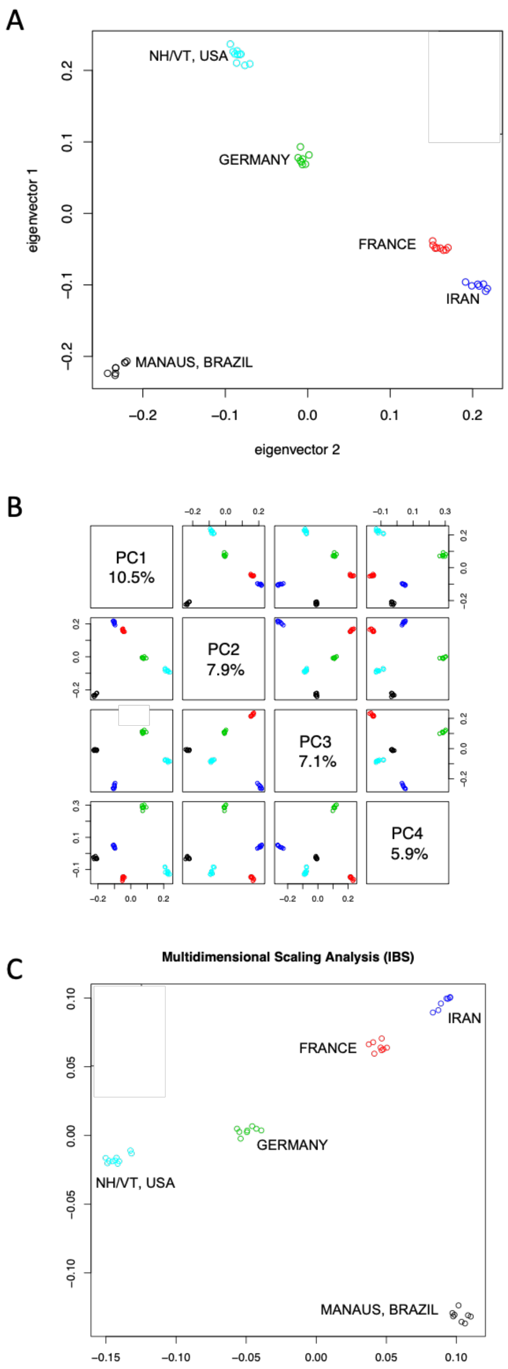


<u>**Figure S11:**</u> **Genomic relationships between five populations of wild house mice.** (**A-B)** Genomic principal component analysis (PCA) across multiple PCs distinguishes mouse populations based on population-of-origin. (**C)** Multidimensional Scaling Analysis (hierarchical clustering) shows similar results to PCA, distinguishing mouse populations based on population-of-origin. <u>Legend</u>: Manaus, Brazil (black); France (red); Germany (green); Iran (dark blue); New Hampshire / Vermont (light blue).

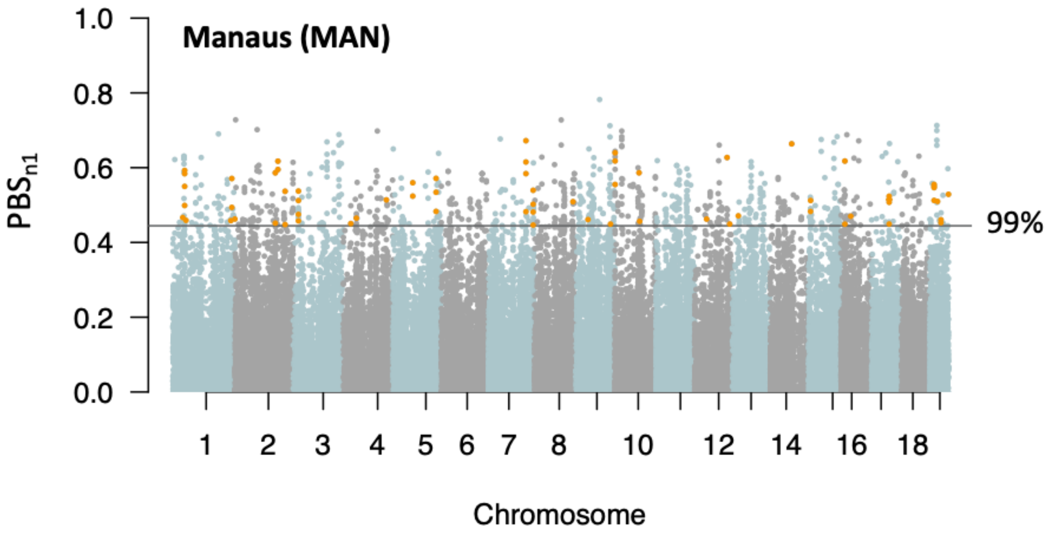


**<u>Figure S12:</u> Autosomal selection scan for the Manaus (MAN) focal population.** Orange points depict genes that exhibit *cis*-regulatory divergence and overlap with *PBSn1* outlier regions.

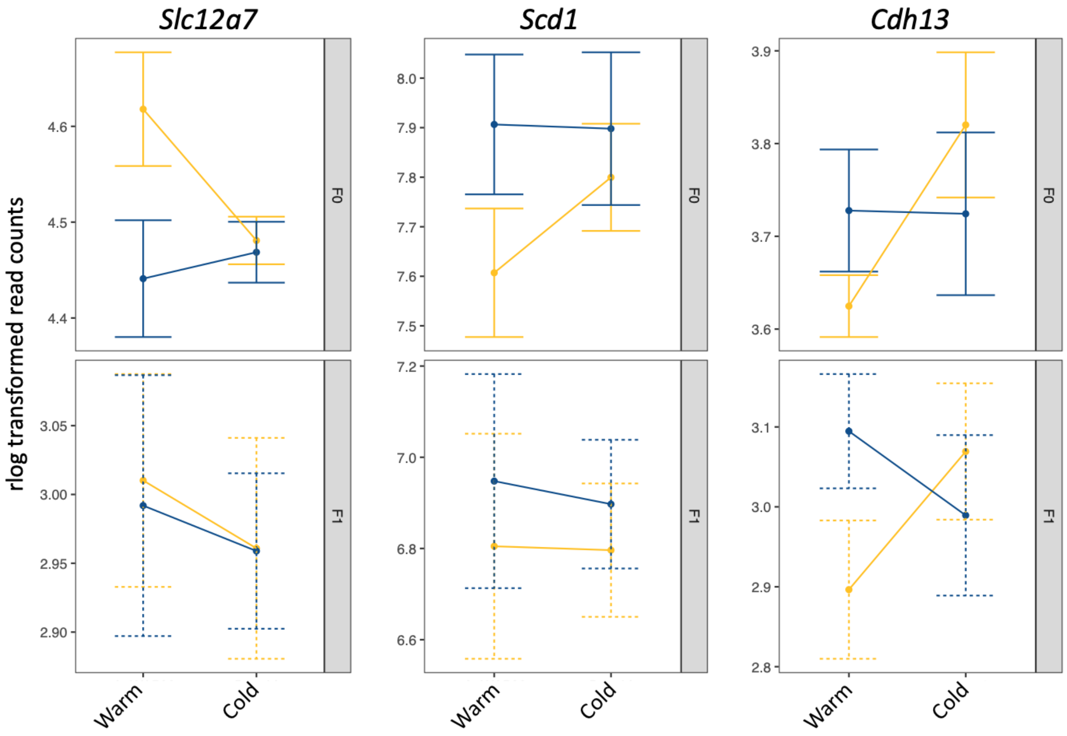


**<u>Figure S13.</u> Expression patterns of three candidate genes showing adaptive plasticity and significant *trans* x environment interaction (*Slc12a7*), *cis* x environment interaction (*Scd1*), and *cis* + *trans* x environment interaction (*Cdh13*)**. Parental expression (F0) and allelic expression (F1) are plotted as regularized log transformed counts.

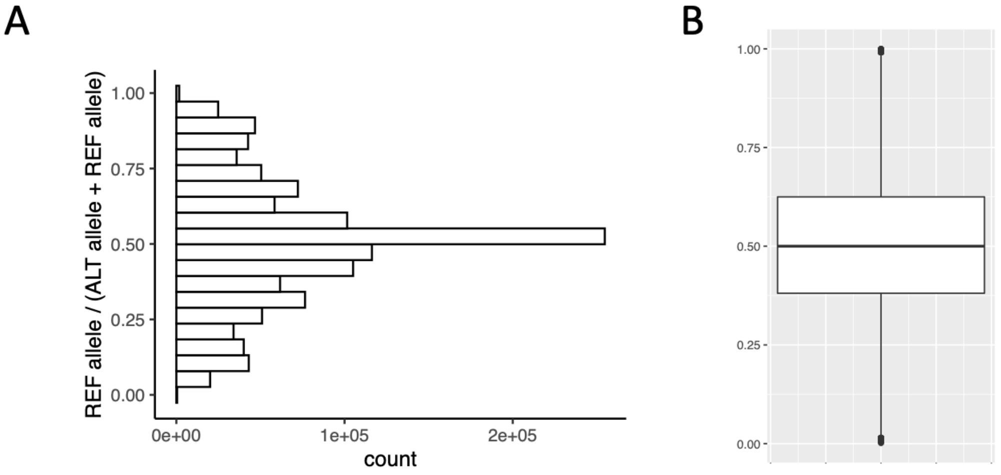


**<u>Figure S14.</u> No evidence for reference mapping bias in allele specific expression analyses. (A)** Distribution of reads overlapping the references vs. alternative allele (REF allele / (ALT allele + REF allele). **(B)** These proportions show a median of 0.5 across samples.

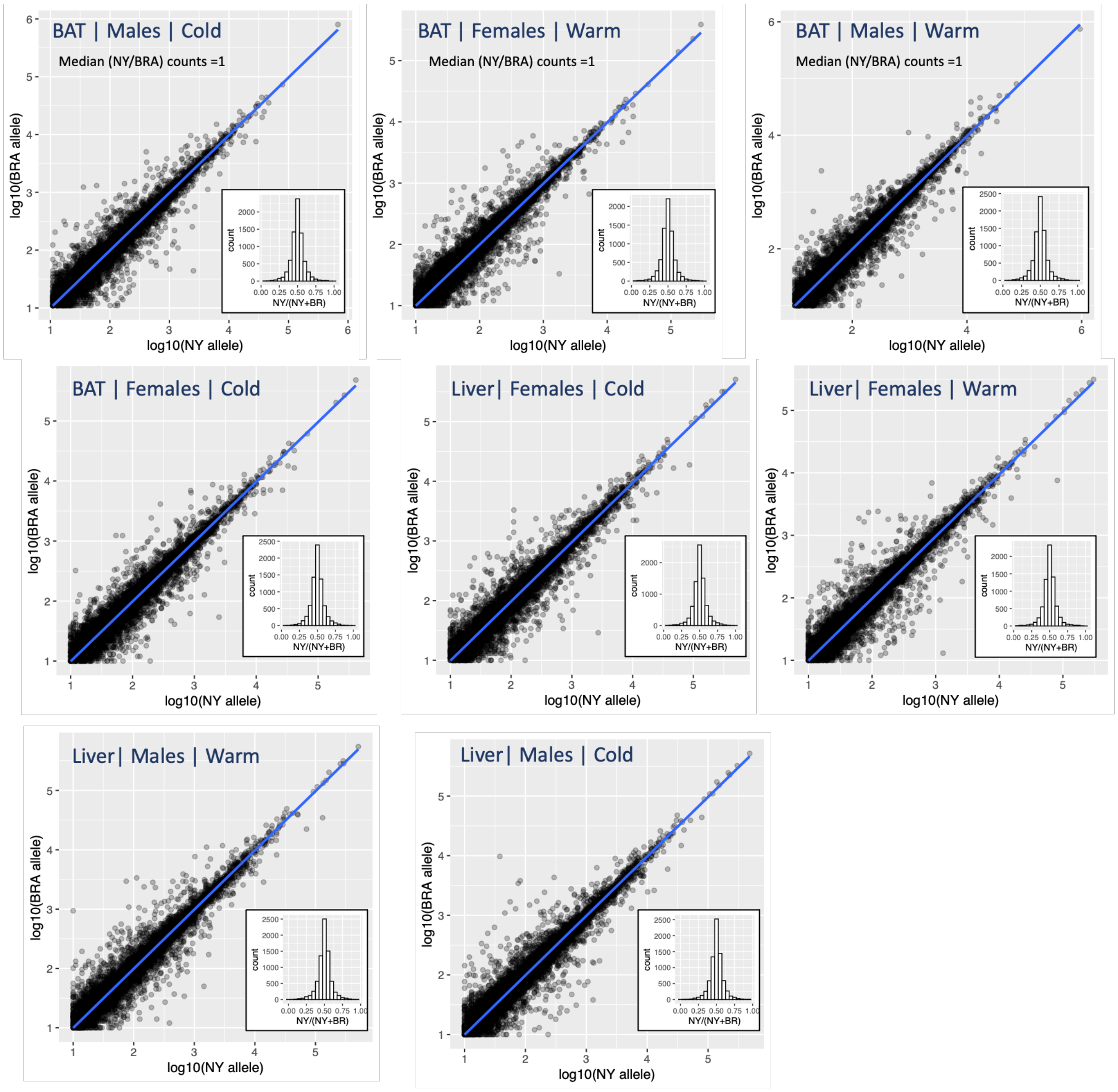


**<u>Figure S15</u>. Further evidence for no reference mapping bias in allele specific expression analyses.** Each plot depicts the number of mapped reads to the BZ and NY allele for each gene in each tissue-type, sex, and environment. For each group, the linear correlation has a slope near 1, indicating little evidence for mapping bias.

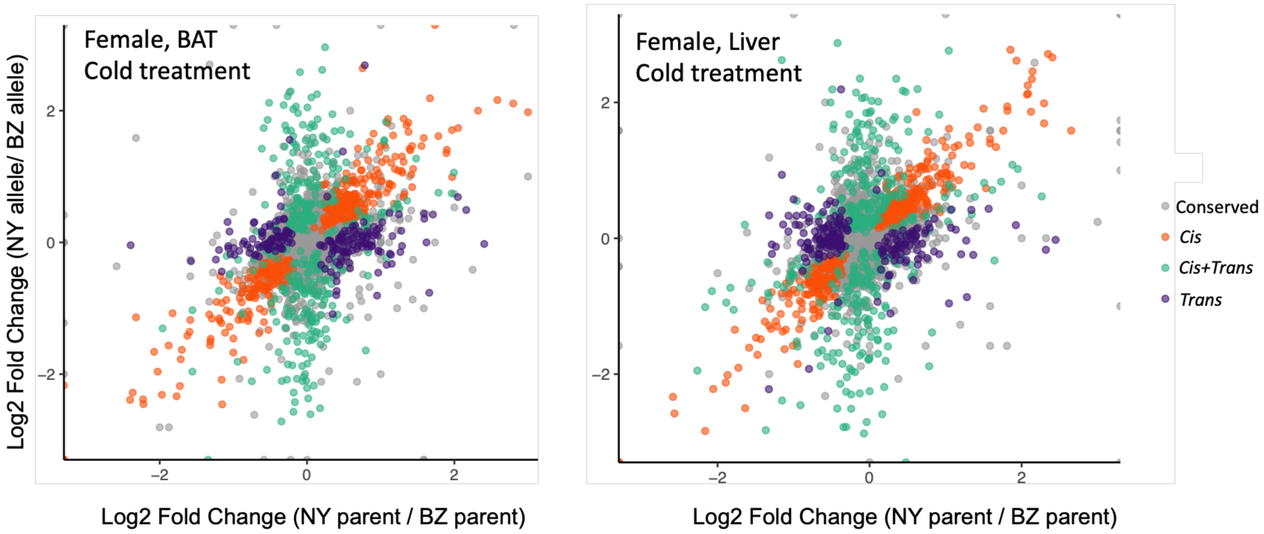


**<u>Figure S16</u>. Categorization of female regulatory divergence.** Points (individual genes) represent log2-fold changes between reads mapping to each allele in the hybrid (BZ allele / NY allele; y-axis) and the reads mapping to each parental line (BZ parent / NY parent; x-axis). Genes are colored based on their inferred regulatory category: orange = *cis*, purple = *trans*, green = *cis&trans*, gray = conserved or ambiguous. Genes categorized as conserved or ambiguous (gray points) constitute roughly 75% of all genes and are centered on the origin and mostly hidden behind other genes.

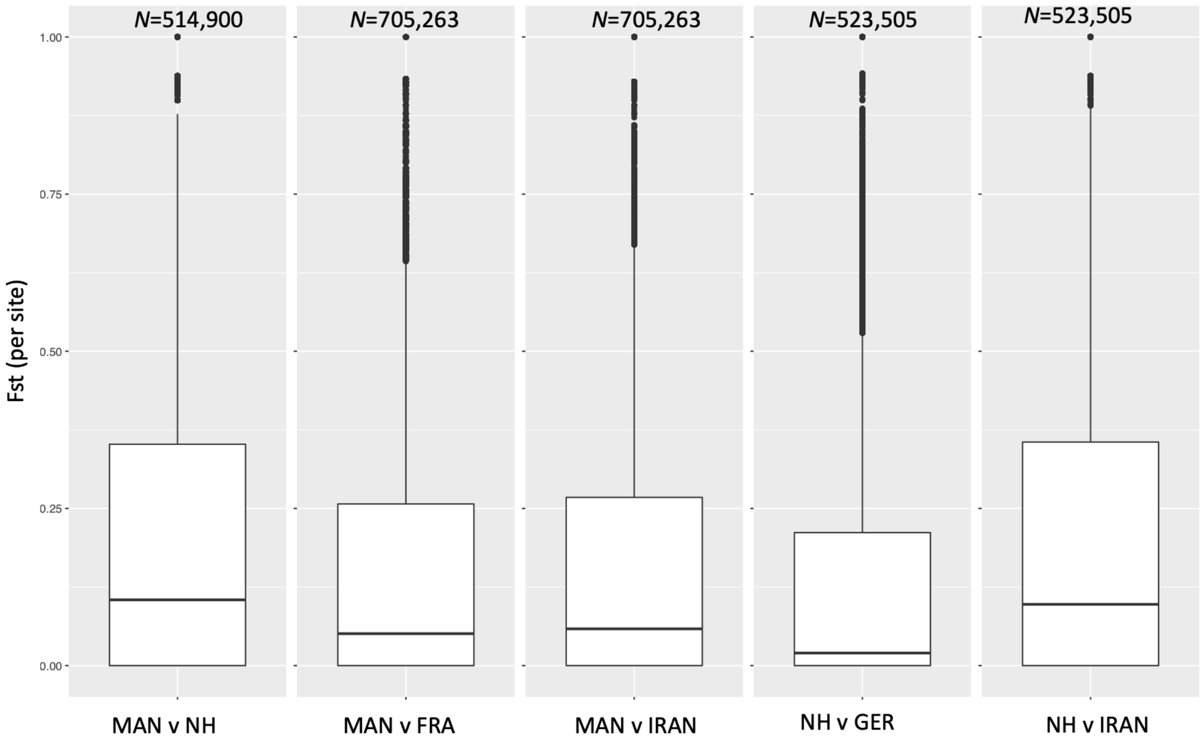


**<u>Figure S17:</u> Pairwise Fst (per site) of *M.m. domesticus* populations used in *PBSn1* analysis.** Number of sites tested (*N*) is depicted above each comparison.

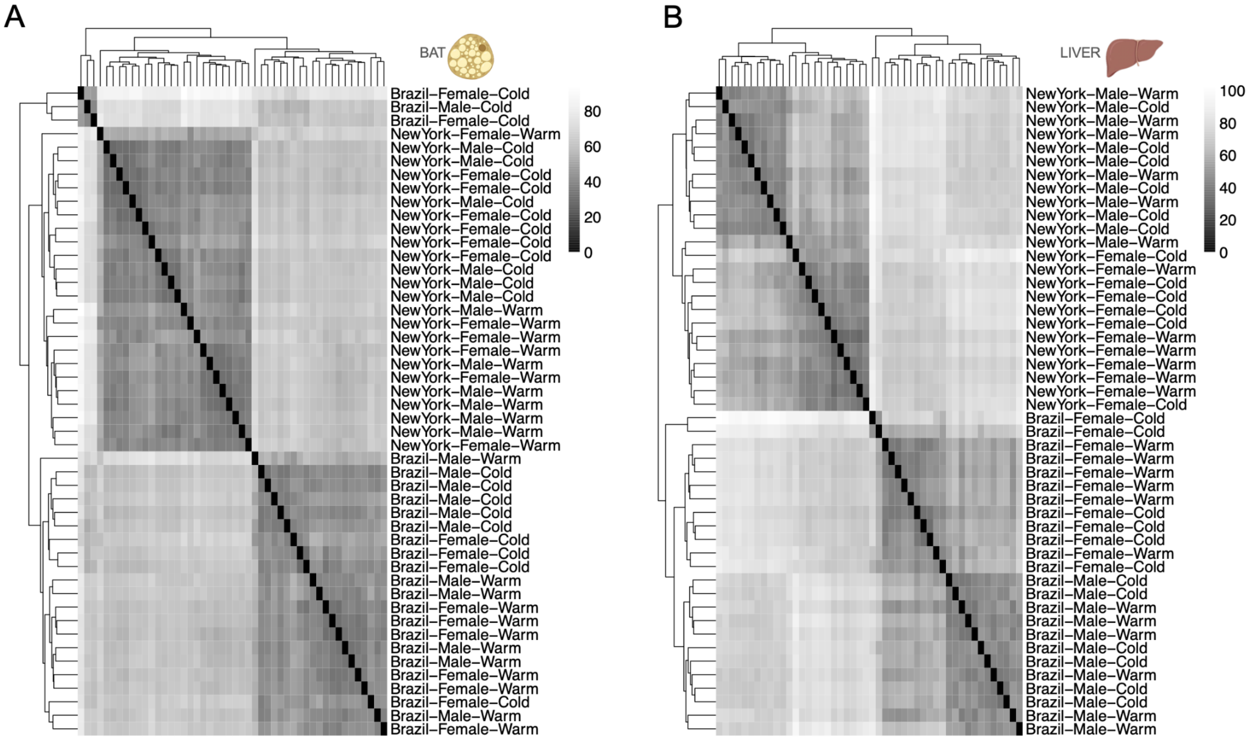


**<u>Figure S18.</u> Hierarchical clustering of sample-to-sample distances in (A) BAT and (B) liver.** For both tissues, samples largely cluster by genotype. In BAT, samples do not cluster by sex.

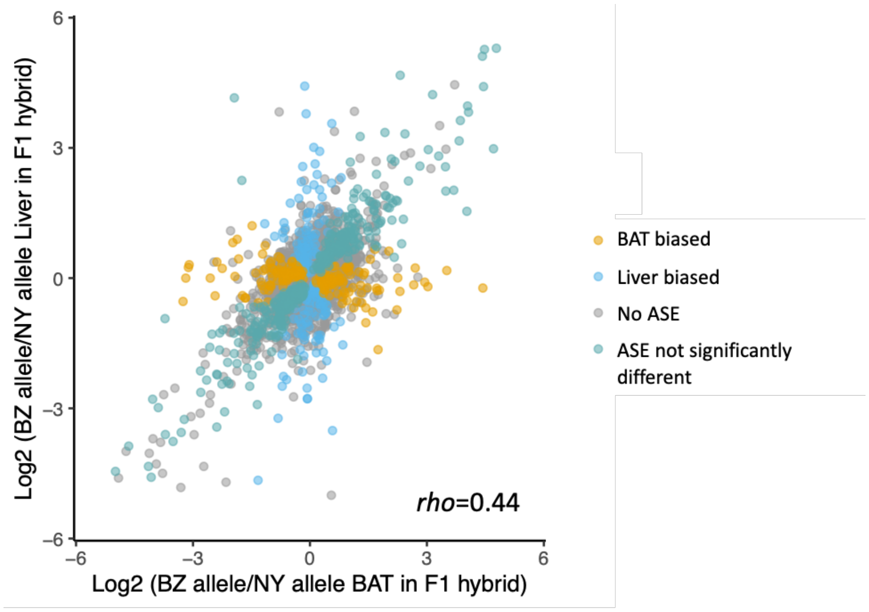


**Figure S19.** **Tissue-dependent allele-specific expression.** Scatterplot comparing the distribution of allelic ratios between BAT and liver. Points (representing individual genes) are colored by binned differential ASE *P*-values on Brazil and New York allele counts in the two tissues. Genes in gray are not genes for which ASE is shared between tissues, but are the background set of all genes tested. Genes in green depict ASE ratios that are not significantly different across tissues, while genes in blue (liver-biased) and orange (BAT-biased) depict tissue-specific ASE. The ratios of BZ to NY read counts are correlated between tissues, suggesting shared regulatory divergence between BAT and liver (Spearman's rho=0.44).

#### Supplemental Tables

**Table S1.** **Results of linear mixed models investigating the effects of sex and genotype (line) on body mass, pelage conductance, and extremity length in house mice.**

| Trait | Factor | $\chi^2$ | DF | *P*-value | Effect size ($\omega^2$) |
|---|---|---|---|---|---|
| Body Mass (g) | | | | | |
| | Line | 28.54 | 1 | < 0.001 | 0.48 |
| | Sex | 15.29 | 1 | <0.001 | 0.31 |
| Pelage Conductance (W $m^{-2}$ °$C^{-1}$) | | | | | |
| | Line | 21.65 | 1 | < 0.001 | 0.35 |
| | Sex | 0.05 | 1 | 0.82 | -0.03 |
| Relative Tail Length (mm/g) | | | | | |
| | Line | 85.97 | 1 | < 0.001 | 0.74 |
| | Sex | 12.12 | 1 | < 0.001 | 0.26 |
| Relative Ear Length (mm/g) | | | | | |
| | Line | 66.67 | 1 | < 0.001 | 0.73 |
| | Sex | 18.55 | 1 | < 0.001 | 0.35 |

**<u>Tables S2.</u> Number of genes exhibiting genotype-by-environment interactions (GxE) across multiple log2 fold change cut-offs.** Analysis (A) refers to GxE patterns depicted in Figures 2B and S3C, while Analysis (B) refers to the relationship between plasticity and divergence depicted in Figures 2C and S3D.

| Analysis | Tissue | Sex | Comparison | all LFC | LFC >= \|0.5\| | LFC >= \|1\| | LFC >= \|1.5\| |
|---|---|---|---|---|---|---|---|
| A | | | | | | | |
| | Liver | | | | | | |
| | | M | | | | | |
| | | | $NY_{warm}$ vs $NY_{cold}$ | 105 | 79 | 33 | 16 |
| | | | $BZ_{warm}$ vs $BZ_{cold}$ | 711 | 281 | 85 | 31 |
| | | F | | | | | |
| | | | $NY_{warm}$ vs $NY_{cold}$ | 98 | 75 | 34 | 18 |
| | | | $BZ_{warm}$ vs $BZ_{cold}$ | 551 | 316 | 168 | 103 |
| | BAT | | | | | | |
| | | M | | | | | |
| | | | $NY_{warm}$ vs $NY_{cold}$ | 485 | 293 | 96 | 39 |
| | | | $BZ_{warm}$ vs $BZ_{cold}$ | 936 | 371 | 99 | 28 |
| | | F | | | | | |
| | | | $NY_{warm}$ vs $NY_{cold}$ | 284 | 170 | 79 | 31 |
| | | | $BZ_{warm}$ vs $BZ_{cold}$ | 1,325 | 694 | 205 | 72 |
| B | | | | | | | |
| | Liver | | | | | | |
| | | M | | | | | |
| | | | same | 296 | 169 | 79 | 46 |
| | | | opposite | 111 | 72 | 34 | 25 |
| | | F | | | | | |
| | | | same | 142 | 107 | 55 | 37 |
| | | | opposite | 56 | 41 | 23 | 15 |
| | BAT | | | | | | |
| | | M | | | | | |
| | | | same | 295 | 199 | 85 | 52 |
| | | | opposite | 203 | 141 | 75 | 40 |
| | | F | | | | | |
| | | | same | 302 | 233 | 138 | 95 |
| | | | opposite | 173 | 125 | 66 | 32 |

**<u>Table S3.</u> Genes with significant *trans* x temperature effects for each tissue.**

| Tissue | Gene | Ensembl Number |
|---|---|---|
| BAT | | |
| | *arl6ip6* | ENSMUSG00000026960 |
| | *hsd11b1* | ENSMUSG00000016194 |
| | *parm1* | ENSMUSG00000034981 |
| | *wnt11* | ENSMUSG00000015957 |
| | *gstt1* | ENSMUSG00000001663 |
| | *cdh13* | ENSMUSG00000031841 |
| | *tmem45b* | ENSMUSG00000041737 |
| | *prpf40a* | ENSMUSG00000061136 |
| | *ccdc3* | ENSMUSG00000026676 |
| | *xylt1* | ENSMUSG00000030657 |
| | *amy1* | ENSMUSG00000074264 |
| | *erbb2* | ENSMUSG00000062312 |
| | *itih5* | ENSMUSG00000025780 |
| | *adamts9* | ENSMUSG00000030022 |
| | *sec1414* | ENSMUSG00000019368 |
| | *slc1414* | ENSMUSG00000017756 |
| | *fam111a* | ENSMUSG00000024691 |
| | *arhgef10* | ENSMUSG00000071176 |
| Liver | | |
| | *hmgcs2* | ENSMUSG00000027875 |

**Table S4. Genes with ASE that co-localize with genomic outliers for NH/VT and are candidates for metabolic adaptation based on mutant phenotypes.**

| Gene | Mutant Phenotypes |
|---|---|
| *col6a1* | body weight and size; metabolism (glucose tolerance) |
| *prkar2b* | body weight; body fat composition; diet-induced obesity; energy expenditure; food intake; body temperature; metabolism (e.g., cholesterol, leptin, triglyceride levels) |
| *col5a3* | body weight; diet-induced obesity; metabolism (e.g., glucose tolerance, insulin levels) |
| *sulf2* | body weight and size |
| *enpp3* | susceptibility to weight loss |
| *smoc1* | body weight and size; body fat; activity level |
| *myo10* | body weight and size |
| *prkdc* | body size and weight gain |
| *stard4* | body weight and size; metabolism (cholesterol level) |
| *impact* | body weight and size; diet-induced obesity; food intake; fat composition; body temperature; metabolism (e.g., glucose level, insulin sensitivity) |
| *lipa* | body weight; body fat composition; food intake; metabolism (e.g., triglyceride, lipid, cholesterol level, insulin resistance) |
| *als2* | body weight |
| *col5a2* | body weight and size |
| *col3a1* | body size |
| *mpzl1* | weight gain |
| *stam* | postnatal growth |
| *bcat2* | body weight; diet-induced obesity; body fat composition; energy expenditure; body temperature; metabolism (e.g., leptin, adiponectin, glucose, insulin levels) |
| *stim1* | postnatal growth |
| *wrn* | susceptibility to age related obesity; body fat composition; metabolism (e.g., triglyceride, cholesterol, insulin levels) |
| *ets1* | body weight; postnatal growth |

| | |
|---|---|
| *myo6* | body size; activity level; metabolism (e.g., glucose, triglyceride level) |
| *tpst1* | body weight |
| *sbno2* | body weight |
| *cc2d2a* | body weight; postnatal growth |
| *b3glct* | body size |
| *plaat3* | susceptibility to weight gain; body fat composition; susceptibility to diet-induced obesity; metabolism (e.g., leptin, adiponectin, glucose levels) |
| *ahnak* | body weight; susceptibility to diet-induced obesity; body fat composition; metabolism (e.g., glucose tolerance) |
| *cog1* | lean body mass |
| *tmem106b* | body size; body fat amount; metabolism (insulin level, glucose tolerance) |
| *vps13a* | body fat amount; activity |
| *wwp1* | activity level; metabolism (glucose level) |
| *dhx29* | activity level; lean body mass; metabolism (glucose tolerance) |
| *sypl* | metabolism (glucose level) |
| *arhgef10* | activity level; metabolism (glucose level) |
| *gpr146* | metabolism (e.g., cholesterol, triglyceride level) |
| *rsad1* | metabolism (e.g., triglyceride level) |
| *pctp* | metabolism (e.g., cholesterol level, lipid homeostasis) |
| *uaca* | metabolism (cholesterol level) |
| *slc46a3* | activity, metabolism (cholesterol level) |
| *cd44* | metabolism (lipid level) |
| *mmd* | body size; activity |
| *zfyve1* | body size |

**<u>Table S5.</u> Number of differentially expressed (DE) genes across parental expression data for each of the four main effects (tissue, sex, genotype, environment) and their interactions using a full parameterized model in DESeq2 (see section 3: "Extended Parental Gene Expression Analyses ").**

| Effect/Interaction | No. DE Genes (FDR < 0.05) |
|---|---|
| Tissue | 13,713 |
| Sex | 1,705 |
| Genotype | 8,480 |
| Environment | 4,446 |
| Tissue:Sex | 1,483 |
| Tissue:Genotype | 5,621 |
| Tissue:Environment | 2,195 |
| Genotype:Sex | 47 |
| Sex:Environment | 1 |
| Genotype:Environment | 126 |
| Tissue:Genotype:Sex | 19 |
| Tissue:Sex:Environment | 2 |
| Tissue:Genotype:Environment | 20 |
| Sex:Genotype:Environment | 35 |
| Tissue:Sex:Genotype:Environment | 0 |

#### SI Dataset S1 (Data_S1.xlsx)

Individual metadata and candidate genes. This file includes:

Sheet 1: Phenotypic metadata for SARA and MANA

Sheet 2: Metadata for RNA-seq samples and associated MVZ accession numbers

Sheet 3: Genes overlapping *PBSn1* outlier blocks in New Hampshire / Vermont (NH/VT)

Sheet 4: Genes overlapping *PBSn1* outlier blocks in Manaus (MAN)

Sheet 5: Genes with ASE overlapping *PBSn1* outliers in MAN and NH/VT

Sheet 6: Phenotype enrichments of outlier genes with ASE in both warm and cold in NH/VT
